# High lineage survivorship across the end-Devonian Mass Extinction suggested by a remarkable new Late Devonian actinopterygian

**DOI:** 10.1101/2021.09.02.458676

**Authors:** Sam Giles, Kara Feilich, Rachel Warnock, Stephanie E. Pierce, Matt Friedman

## Abstract

A mass extinction at the end of the Devonian is thought to have had a major influence on the evolution of actinopterygians (ray-finned fishes), which comprise half of living vertebrates. This extinction appears to have acted as a bottleneck, paring the early diversity of the group to a handful of survivors. Coupled with increases in taxonomic and morphological diversity in the Carboniferous, this contributes to a model of explosive post-extinction radiation. However, most actinopterygians from within a ~20-million-year (Myr) window surrounding the extinction remain poorly known, contributing to uncertainty about these patterns. An exceptionally preserved fossil of a diminutive fish from 7 Myr before the extinction reveals unexpected anatomical features that suggest a very different story. This new fossil nests within a clade of post-Devonian species and, in an expanded phylogenetic analysis, draws multiple lineages of Carboniferous actinopterygians into the Devonian. This suggests cryptic but extensive lineage diversification in the latest Devonian, followed by more conspicuous feeding and locomotor structure diversification in the Carboniferous. Our revised model matches more complex patterns of divergence, survival, and diversification around the Devonian-Carboniferous boundary in other vertebrate clades. It also fundamentally recalibrates the onset of diversification early in the history of this major radiation.

## Main

The distinction between Devonian and Carboniferous vertebrate faunas has long been apparent^1,2^. Debate on the underlying cause of this faunal turnover focuses on whether it reflects a sudden transition stemming from a mass extinction event followed by explosive evolutionary recovery^3,4^, or a more gradual shift obscured by incomplete or understudied paleontological evidence^5,6^. Taken at face value, the fossil record of actinopterygians (ray-finned fishes) appears to provide strong evidence for the former hypothesis. Actinopterygians persisted at low diversity^3^ and abundance^7,8^ throughout the Devonian, but assumed their role as the dominant group of aquatic vertebrates in the early Carboniferous^3^. Two important patterns of phenotypic change are associated with this taxonomic shift. First, Carboniferous actinopterygians show a significantly greater repertoire of body and skull shapes than their Devonian predecessors^9^, including feeding ^10,11^ and locomotory^12^ innovations. Second, there are indications of bodysize reduction in actinopterygians and other vertebrate lineages at this time, interpreted as a protracted ‘Lilliput effect’ ^13^.

Unstable phylogenetic relationships and a rudimentary understanding of latest Devonian and earliest Carboniferous actinopterygians complicates interpretation of these patterns. Consensus views on phylogeny indicate only one or two actinopterygian lineages crossed the Devonian-Carboniferous boundary^14–16^ (but see^17^), suggesting that prolific diversity in the early Carboniferous arose through an explosive evolutionary radiation seeded by a handful of surviving groups, a mechanism historically proposed for famous examples like placental mammals^18^. Recent analysis suggests an additional Devonian lineage persisting into the early Carboniferous alongside a single radiation containing all Carboniferous and younger taxa^19^. These models contrast with the emerging picture for other vertebrate survivors of the end-Devonian extinction: divergence of multiple lineages in the Late Devonian, followed by substantial ecomorphological radiation in the early Carboniferous. Inadequate taxonomic sampling in early actinopterygian phylogenies hinders meaningful discrimination among these models. Most well-known taxa occur millions of years before^20^ or after^15,21,22^ the Devonian-Carboniferous boundary. Taxa from deposits more proximate to the extinction event ^23–26^ are rarely included in systematic analyses because of a limited understanding of their anatomy stemming from their small size^25^, incomplete preservation^26^, or both. However, fossils from the latest Devonian^27^ or earliest Carboniferous ^19,28^ hint at a radically different interpretation of divergences before, survivorship across, and diversification after the end-Devonian extinction.

Here we use micro-computed tomography to provide detailed anatomical information for a new genus and species of actinopterygian from the Late Devonian (middle Famennian; ca. 367 Ma) of the Appalachian Basin of the eastern United States. Previously referred to the problematic genus *Rhadinichthys*^29^, it bears an unexpected combination of derived features previously known only in Carboniferous and younger actinopterygians. This taxon prompts a reconsideration of trait evolution and tree topology that together provide evidence for prolific radiation of ray-finned fishes before the end-Devonian mass extinction. This new perspective on ray-finned fish evolution complements evidence in other groups for more complex patterns of evolution through end-Devonian ecological crises^30–35^.

## Systematic Paleontology

*[redacted from preprint]*

### Diagnosis

Actinopterygian characterized by the following combination of characters: ornament comprising broad ridges incised with narrow grooves; dermosphenotic with well-developed posterior ramus; three suborbitals; ramifying tubules of infraorbital canals in jugal; median perforation in aortic canal of basicranium; two accessory vomers; dermal basipterygoid process; dermohyal unfused to imperforate hyomandibula.

### Holotype

MCZ VPF 5114, a nearly complete fish preserved in part and counterpart, missing snout, anterior portion of lower jaw, and caudal fin (Supplementary Figs. 1–2).

### Horizon and locality

“Chemung” of Warren, Pennsylvania, USA^29^. The Chemung of earlier workers is a facies rather than formal stratigraphic unit, represented in the Chadakoin and Canadaway formations in western Pennsylvania^36,37^. Only the younger of these, the Chadakoin Formation, is present in the collection locality^36^. The Chadakoin Formation lies within the *Palmatolepis marginifera* Conodont Zone (Kirchgasser), Goniatite Zone II-G^38^, and Fa2c subdivision of Spore Zone GF^39^, indicating a mid-Famennian age, providing a constraint of 367.9-367.2 Ma for MCZ VPF 5114^40^. Other fishes reported from the Chadakoin Formation include arthrodire placoderms, chondrichthyans, lungfishes, and porolepiforms^29,41^.

### Remarks

Eastman^29^ referred MCZ VPF 5114 to the problematic genus *Rhadinichthys*, leaving it in open nomenclature as *Rhadinichthys* sp. Species assigned to *Rhadinichthys* do not form a clade^14^, and the holotype of the type species, *R. ornatissmimus*, is poorly preserved (National Museum of Scotland NMS G.1878.18.7). There is no evidence linking MCZ VPF 5114 to this taxon to the exclusion of other early actinopterygians. The small size and incomplete endoskeletal ossification of this specimen may indicate that it is a juvenile.

## Results

### Skull roof

The skull roof comprises frontals, parietals, supratemporals and intertemporals (Fig. 1A, Supplementary Figs. 3–5). The frontal, which is pierced by the pineal foramen, is the largest bone of the skull roof. A midline suture is not visible in tomograms, and it is unclear whether the frontals were tightly joined or fused as a single plate. A notch on the lateral margin of the frontal accommodates the supratemporal. The supraorbital canal extends through the frontal and into the parietal. This canal, housed on the ventral surface of the bone, is fully floored with bone anteriorly but is open ventrally posterior to the level of the pineal foramen. The quadrate parietal bears posterior and middle pit-lines, and a short posterior extension of the supraorbital canal appears as a faint ridge on the outer surface of the bone with a matching groove on its ventral surface. A modest intertemporal lies anterior to the suture between the parietal and frontal. It carries a short segment of the infraorbital canal along its thickened ventrolateral margin. The lenticular supratemporal is very long, and the infraorbital canal in a ridge along its lateral margin. The lateral margin of the supratemporal abuts the preoperculum, and posterior to this contact a branch of the infraorbital canal extends toward the preopercular canal to form a jugal canal.

**Figure 1.**
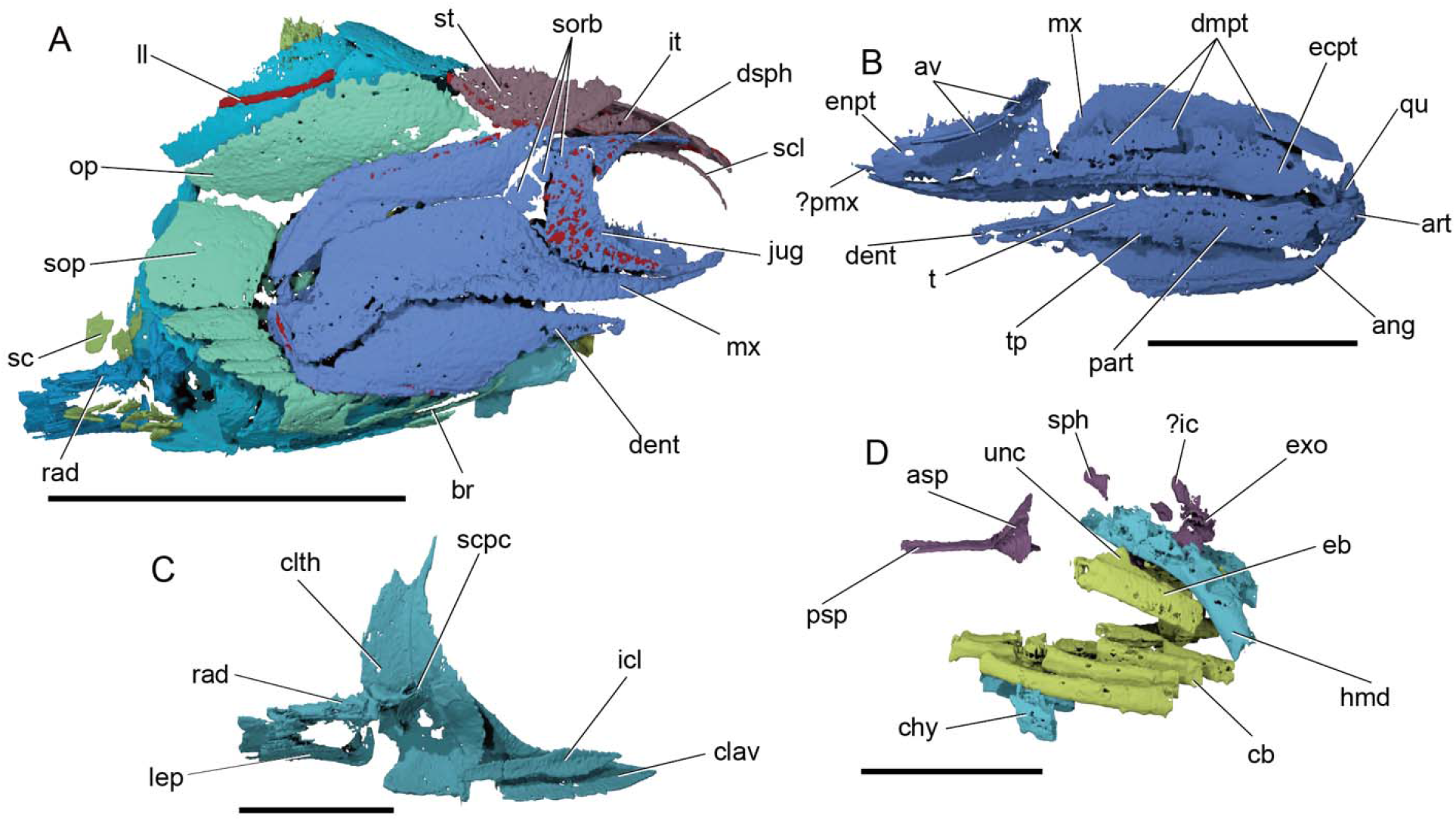
Anatomy of MCZ VPF 5114 [new taxon name redacted from preprint]. Renders of (A) right lateral view of specimen; (B) medial view of right upper and lower jaws; (C) medial view of left shoulder girdle and pectoral fin, and (D) left lateral view of gill skeleton, hyoid arch and braincase. Colour coding of skeleton: blue, cheek and jaw; purple, skull roof and sclerotic ossicle; pink, braincase; dark green, hyomandibula; light green, operculogular system; turquoise, shoulder girdle; yellow, gill skeleton. Scale bar in (A): 10 mm, in (B)-(D): 5 mm. Abbreviations: ang, angular; art, articular; asp, ascending process of parasphenoid; av, accessory vomer; br, branchiostegal; cb, ceratobranchial; chy, ceratohyal; clav, clavicle; dent, dentary; dmpt, dermatopterygoid; dsph, dermosphenotic; eb, epibranchial; ecpt, ectopterygoid; enpt, entopterygoid; exo, exoccipital; hmd, hyomandibula; ?ic, possible intercalar; icl, interclavicle; it, intertemporal; jug, jugal; lep, lepidotrichia; ll, lateral line; mx, maxilla; op, operculum; part, prearticular and coronoid; ?pmx, possible premaxilla; psp, parasphenoid; qu, quadrate; rad, radials; sc, scale; scl, sclerotic ossicle; scpc, scapulocoracoid; sop, suboperculum; sorb, suborbitals; sph, sphenotic ossification; st, supratemporal; t, teeth; tp, toothplate; unc, uncinate process.

### Cheek

The cheek comprises a dermosphenotic, jugal, lacrimal, three suborbitals, and preoperculum (Figs. 1A, 2A, Supplementary Figs. 3–5). The dermosphenotic is ‘T’-shaped, with long anterior and posterior limbs and a short ventral limb. The infraorbital canal extends through the ventral limb of the bone into the jugal, and a branch in the anterior limb of the dermosphenotic approaches but does not connect with the supraorbital canal of the frontal. Within the jugal, the canal is offset from the orbital margin, and at least 14 branches radiate into its posterior portion. The lacrimal is formed as a tube of bone surrounding the infraorbital canal. Three suborbitals lie between the dorsal limb of the jugal and the anterior margin of the preoperculum. They are arranged in a vertical series, with the middle suborbital smaller than the dorsal and ventral ones. The preoperculum, which bears two pit lines, is boomerang-shaped, with an anterior limb broader than the ventral limb. The angle between these limbs bears a broad overlap area for the maxilla. The preopercular canal extends in a thickened ridge on the inner surface of the bone, near its dorsal margin. The presence of a quadratojugal is uncertain.

**Figure 2.**
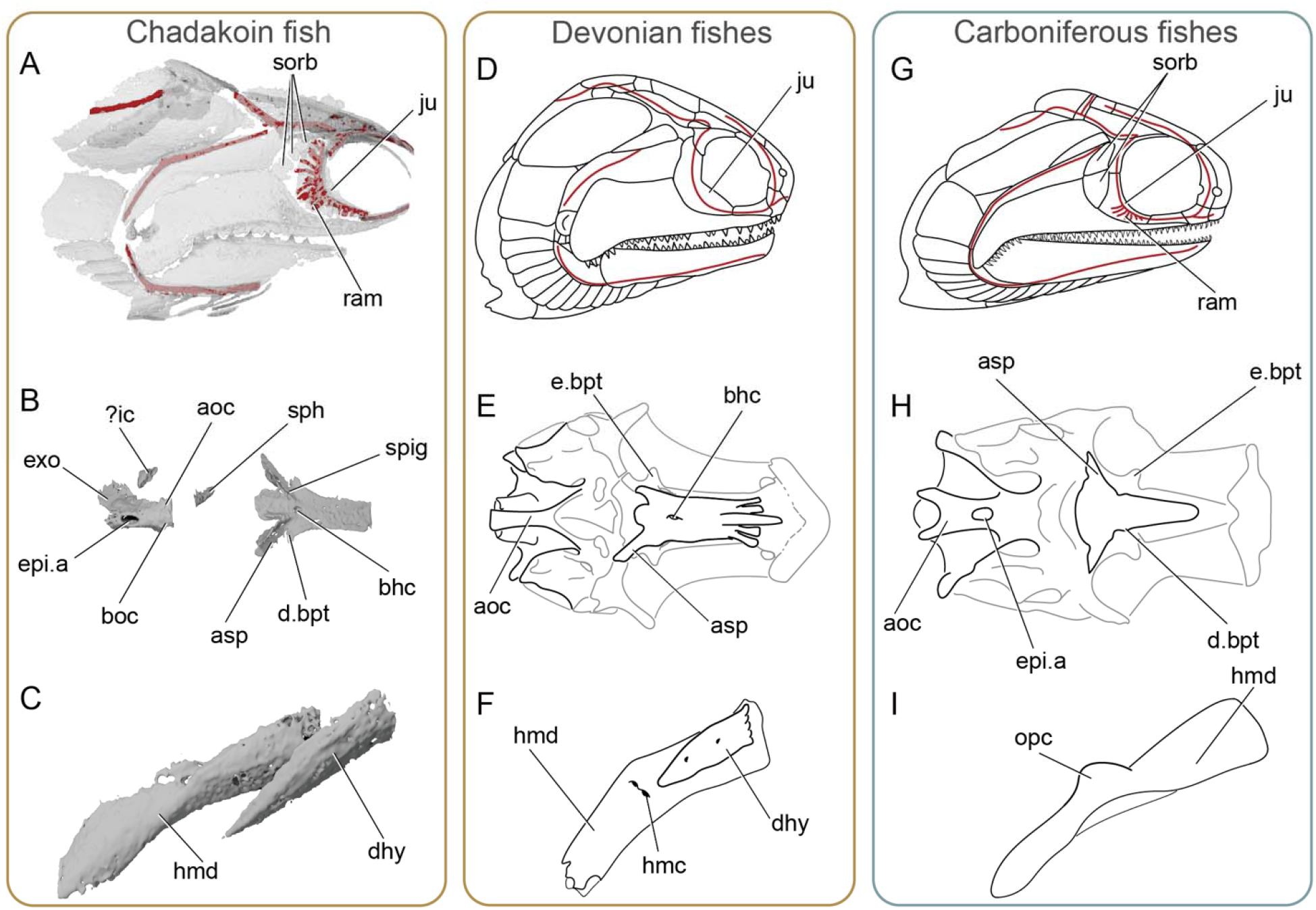
Comparative anatomy of MCZ VPF 5114 [new taxon name redacted from preprint] with Devonian and Carboniferous actinopterygians. Renders of MCZ VPF 5114 [new taxon name redacted from preprint] showing (A) right cheek in lateral view with sensory canals visible, (B) braincase in ventral view, and (C) right hyomandibula in lateral view. Interpretive drawings of *Mimipiscis* showing (D) right cheek in lateral view with sensory canals visible and (E) braincase in ventral view. Interpretive drawing of (F) *Moythomasia durgaringa* showing right hyomandibula in lateral view. Interpretive drawings of (G) *Cosmoptychius striatus* showing right cheek in lateral view with sensory canals visible, (H) *Lawrenciella schaefferi* showing braincase in ventral view, and (I) an indeterminate Kasimovian actinopterygian showing right hyomandibula in lateral view. Parts of the braincase are masked out in *Mimipiscis* and *Lawrenciella* to facilitate comparison with of MCZ VPF 5114 [new taxon name redacted from preprint]. (D)-(F) redrawn from Gardiner ^20^, (H) redrawn from Gardiner ^68^, (I) redrawn from Hamel and Poplin ^43^, and (J) redrawn from Poplin^22^. Not to scale. Abbreviations: aoc, aortic canal; asp, ascending process of parasphenoid; bhc, buccohypophysial canal; boc, basioccipital; d.bpt, dermal basipterygoid process; dhy, dermohyal; e.bpt, endoskeletal basipterygoid process; epi.a, epibranchial artery; exo, exoccipital; hmc, hyomandibular canal; hmd, hyomandibula; ?ic, possible intercalar; ju, jugal; opc, opercular process; ram, ramifying tubules of infraorbital sensory canal; sorb, suborbitals; sph, sphenotic ossification; spig, spiracular groove;

### Upper jaw and palate

Preserved components of the upper jaw include the maxilla, a putative premaxilla, the quadrate and the dermal palatal bones (Fig. 1A,B, Supplementary Figs. 3–6).The maxilla is cleaver-shaped and its posterior margin is very thin and difficult to discern. A posteroventral extension of the posterior plate overlaps the lower jaw. A narrow shelf extends along the medial surface of the bone, beginning at the tip of the suborbital limb and continuing to the lower jaw overlap. Ornament on the maxilla comprises a series of broad, flat ridges that are only narrowly separated from each other, resembling a plateau of enamel separated by narrow grooves. Maxillary teeth are arranged in a row along the ventral margin of the bone, but only sockets are preserved. Marginal teeth are not visible, but this may represent a limit to scan resolution. An elongate, narrow strip of dermal tooth-bearing bone medial to the maxilla may represent a displaced premaxilla.

The endoskeletal portion of the palate is largely unmineralized. Only the quadrate is well-ossified, with two prominent condyles. The dermal ossifications of the palate are preserved as separate bones, but divisions between them are not always clear. The ectopterygoid is long and ovoid, with an irregular anterior margin but narrowing to a point near the quadrate posteriorly. Dorsally, it is bordered by at least three separately ossified dermatopterygoids, although a gap between two of these suggests four were present in total. Ventral to the entopterygoid, the demopalatines presumably form a long narrow extension continuous with the ectopterygoid, although no sutures between them are visible. The entopterygoid, which is roughly rectangular, lies dorsal to the anterior half of the demopalatine series.

Two accessory vomers on each side of the skull are closely associated with the parasphenoid. The anterior accessory vomer is approximately twice the length of the posterior one, and tapers slightly anteriorly. It is confined anterior to the basipterygoid process and broadly overlaps the parasphenoid, while the posterior accessory vomer traces the spiracular groove on the ascending process of the parasphenoid.

### Lower jaw

The lower jaw is incomplete anteriorly but comprises dermal components of the outer and inner surface and the articular (Fig. 1A,B, Supplementary Figs. 3–5). The external surface of the lower jaw includes a dentary and at least one postdentary bone; the bone in the posterodorsal portion of the lower jaw is very thin, but the dentary-postdentary suture is visible externally. Ornament on the dentary resembles that of the maxilla. The mandibular canal traverses the ventral margin of the dentary and the posteroventral margin of the postdentary. A single row of seven large conical teeth is present along the dorsal margin of the dentary, plus at least seven empty sockets. No marginal dentition is visible, although this may be an artefact of scan resolution.

The mesial surface of the lower jaw is covered by dermal bone, probably representing the prearticular and coronoids, although divisions between bones are indistinct. This complex defines the mesial margin of the adductor fossa posteriorly. A longitudinal ridge is present on the dorsal surface of these dermal bones. No distinct teeth are visible, but modest crenulation of this ridge suggests that small denticles were present. Three irregularly shaped toothplates are applied to the exposed mesial surface of the prearticular/coronoid complex. Narrow gaps separate individual toothplates, which collectively form a large elliptical composite plate.

Only the articular region of Meckel’s cartilage is ossified. It bears concave mesial and lateral facets that match the convex condyles on the quadrate.

### Neurocranium and parasphenoid

The braincase is only partially ossified, and is floored by a short parasphenoid (Fig. 1D, 2B, Supplementary Figs. 3–4,6). Mineralization of the neurocranium is largely restricted to the otic and occipital region, comprising a basioccipital plate; dorsal, paired ossifications corresponding to the exoccipital components of the occipital arch; paired, nodular ossifications anterior to the exoccipitals possibly representing intercalars, and a small ossification posterodorsal to the orbit in the sphenotic region. A well-developed aortic canal, pierced by a prominent midline foramen for the efferent artery, extends the length of the basioccipital ossification. The exoccipital ossifications preserve a single foramen on their lateral surfaces, possibly for a spinoccipital nerve.

The parasphenoid comprises a plank-like anterior corpus and a short posterior extension with a rounded margin. Because of poor endocranial mineralization, it is unclear if it bridged a ventral fissure. In crosssection, the anterior corpus is concave ventrally and convex dorsally. A broad, rounded ridge extends along the dorsal midline of the anterior half of the corpus. A denticulated ridge on the ventral surface of the corpus was exposed between the accessory vomers. Three major structures are present near the junction between the anterior corpus and posterior extension of the parasphenoid: a prominent midline buccohypophysial foramen, modest dermal basipterygoid processes, and long ascending processes. Pronounced spiracular grooves extend along the lateral surfaces of the ascending processes. Deep notches mark the posterior intersection of the ascending processes and the body of the parasphenoid, and a parabasal canal is present.

### Hyoid arch and associated dermal bones

Endoskeletal ossifications of the hyoid arch include the hyomandibula and ceratohyal, both of which are incompletely mineralized (Fig. 1D, 2C, Supplementary Figs. 3–6). These are joined by a large dermohyal. The imperforate hyomandibula has a distinct elbow marking the junction of its dorsal and ventral limbs, but no opercular process is present. A groove on the dorsomedial surface of the upper limb marks the course of the hyomandibular branch of the facial nerve. Mineralization of the hyomandibula is incomplete proximally and distally. The plate-like ceratohyal lacks a medial constriction and bears a lateral groove for the afferent hyoid artery. The proximal and distal ends of the bone are unmineralized. Only the right ceratohyal is completely enclosed in rock, with the left bone partially exposed on the surface. The triangular dermohyal is longer than the preserved dorsal shank of the hyomandibula, and the two are unfused.

### Branchial arches

Mineralization of the gill skeleton is generally poor throughout (Fig. 1D, 2D, Supplementary Figs. 3–6). The gill skeleton has four rod-like ceratobranchials bearing a longitudinal ventral groove, with a poorly mineralized and narrow fifth arch visible in the scan but beyond the limit of segmentation. Paired bones representing hypobranchial 1 lie at the anterior of the gill skeleton, displaced from life position. Hypobranchial 1 is one third the length of ceratobranchial 1, and has a ventral process at its proximal end. The basibranchial is largely unossified, but a small segment bearing paired swellings is preserved anterior to the second ceratobranchials. Two epibranchials are mineralized on each side of the skull. Each bears a dorsal longitudinal groove and an uncinate process. A single pharyngobranchial is preserved on each side of the specimen, in articulation with the basioccipital.

### Operculogular series

The operculogular series comprises an operculum, suboperculum, and branchiostegals (Fig 1A, Supplementary Figs. 3–4). The rhombic operculum has a long diagonal axis and small prong at its anterior corner. Ridges ornament the external surface of the bone. The quadrangular suboperculum is half the size of the operculum. Around 13 splint-shaped branchiostegals are preserved, the most posterior of which is the deepest. As the anteriormost branchiostegals are exposed on the surface, it is not possible to determine their exact number or identify any gular plates.

### Pectoral girdle and fin

The dermal shoulder girdle includes extrascapulars, a posttemporal, presupracleithrum, supracleithrum, cleithrum, and clavicle (Fig. 1A,C,D, Supplementary Figs. 3–4,7). As only one half of the skull roof is preserved, the exact number of extrascapulars is unknown, but it seems likely that only one bone was present per side. There is no trace of a medial branch of the lateral line canal. The leaf-shaped posttemporal makes a broad contact with the extrascapular(s), and overlaps the presupracleithrum and supracleithrum. A pronounced groove for the lateral line is visible on the supracleithrum. The cleithrum is broad, with both a laterally-directed and medially-directed face separated medially by a deep ridge. Anterior to the cleithra are paired, strongly curved clavicles. Both clavicle and cleithra are ornamented with long ridges. The anterodorsal portion of the clavicle is developed as a long, narrow spine. The interclavicle is elongate and arrow-shaped, and only a narrow ridge on its ventral surface would have been exposed between the ventral faces of the clavicles. At least three proximate lepidotrichs are fused together at the leading edge of the fin, although they do not appear to embrace the propterygium. The remaining lepidotrichs are evenly segmented and bifurcate distally. Fringing fulcra are absent.

The endoskeletal shoulder girdle and radials are well ossified (Fig. 1C, Supplementary Figs. 3–4,7). The scapulocoracoid is present as several separate ossifications. The largest portion is a curved, horizontal ossification, the lateral face of which abuts the cleithrum and the posterior face of which contacts the propterygium. The remaining radials were presumably supported by a cartilaginous. Three flat, curved plates, each smaller and narrower than the horizontal ossification, comprise the remainder of the scapulocoracoid, but do not appear to be in life position. Three stout, rod-like radials are present between the metapterygium and propterygium. The propterygium is complex in shape, more than twice the size of any other radial and pierced by the propterygial canal. The metapterygium is longer than the remaining radials and supports one preaxial radial.

Little of the remaining fins can be described. Basal scutes are absent. The anal fin bears fringing fulcra, and pelvic fins are present.

### Scales

The scales are rhombic, with an internal ridge and peg-and-socket articulation (Supplementary Figs. 2–4). Ornament comprises broad flat ridges of enamel incised with roughly parallel narrow grooves.

## Discussion

### Anatomical comparisons and phylogenetic placement

Our data reveal internal anatomy unanticipated in a Devonian actinopterygian (Fig. 2A-C). Most prominent are a midline perforation in the aortic canal and dermal basipterygoid processes, previously known only in post-Devonian taxa^22,42–45^. This is joined by features such as suborbitals and multiple rami of the infraorbital canal in the jugal that are only sporadically reported in Devonian species^46,47^ (Fig. 2D-F) but overwhelmingly present in younger taxa (Fig. 2H-J). MCZ VPF 5114 [new taxon name redacted from preprint] is small, but not exceptionally so (total length ~50 mm). While some aspects of its anatomy are consistent with an earlier ontogenetic stage (e.g., limited endoskeletal ossification), comparison with development in extant species indicates that the features that ally it with Carboniferous actinopterygians are not transient juvenile features^48,49^.

This qualitative assessment is corroborated by formal phylogenetic analysis of a published matrix^50^ expanded to incorporate additional Famennian and Tournaisian actinopterygians. Our parsimony (Supplementary Fig. S8) and Bayesian analyses (Supplementary Fig. S9) recover MCZ VPF 5114 [new taxon name redacted from preprint] as a stem actinopterygian in an immediate clade with two Mississippian taxa. In common with previous analyses^16,50^, resolution is poor across the Bayesian tree. In our parsimony analysis, MCZ VPF 5114 [new taxon name redacted from preprint] and its sister taxa are recovered in a broader clade of Carboniferous–Triassic and younger ray-finned fishes. The placement of MCZ VPF 5114 [new taxon name redacted from preprint] as nested within a Carboniferous radiation is supported by characters relating to the dermal and endoskeleton (Dataset S5).

We also recover multiple clades containing both Devonian and stratigraphically younger forms. This contrasts with the majority of recent phylogenetic hypotheses, which generally recover all post-Devonian taxa in just one^16,50^ or two^14,15,20,51^ clades; earlier analyses suggesting more substantial diversification outside of the actinopterygian crown ^17,52^ are likely biased by methodological and sampling issues^53^. MCZ VPF 5114 [new taxon name redacted from preprint] establishes that key anatomical features previously thought restricted to Carboniferous and younger taxa arose in the Devonian, and the nested phylogenetic position of MCZ VPF 5114 [new taxon name redacted from preprint] and other Late Devonian taxa provides explicit support for the hypothesis that numerous actinopterygian lineages crossed the end-Devonian boundary.

### Models of early actinopterygian diversification

Timing of divergences among early actinopterygians in past studies is either taken at face-value from the fossil record ^3,13^ or depicted graphically in trees ^19,54,55^ timescaled using naïve *a posteriori* approaches^56^. Prevailing hypotheses using these approaches posit that a minimal number of lineages crossed the end-Devonian boundary, with explosive diversification occurring in the early Carboniferous (Fig. 3A,B). Even without formal analysis of divergence times, our placement of MCZ VPF 5114 [new taxon name redacted from preprint] instead supports another model: lineage diversification in the Late Devonian that led directly to the post-extinction radiation (Fig. 3C). A fossil birth-death analysis^57^ provides a more explicit timescale for early actinopterygian evolution (Fig. 3D; Supplementary Fig. 10; Dataset S4), and places the origin of crown actinopterygians in the Late Devonian or earliest Carboniferous (mean: 364.4 Myr; 95% highest posterior density: 379.1 Myr–351.6 Myr). The maximum clade credibility tree suggests that at least ten lineages—an order of magnitude more than inferred by most recent analyses^16,50^—persisted into the Carboniferous, indicating substantial and hitherto cryptic diversification before the end-Devonian extinction.

**Figure 3.**
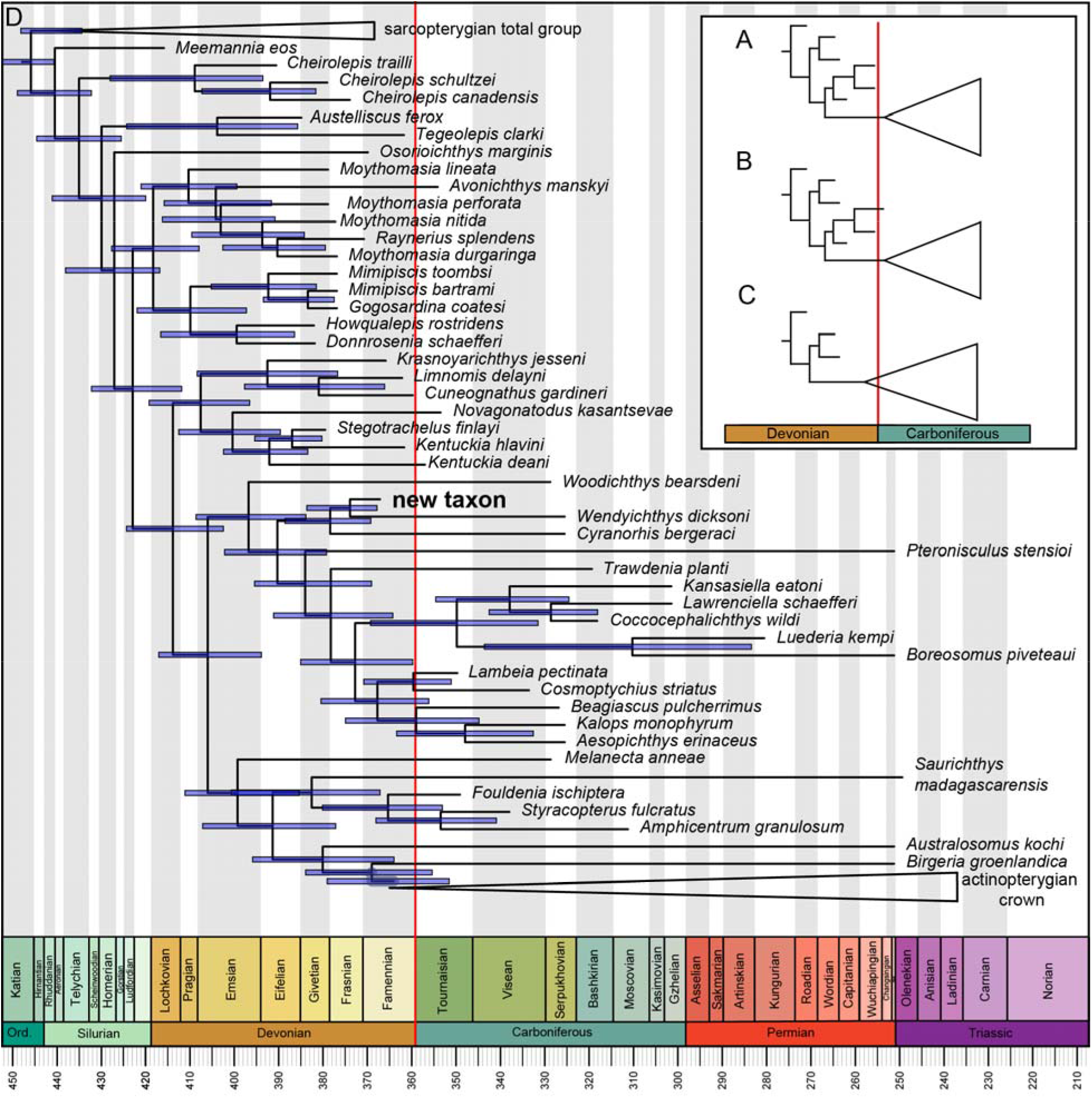
Hypotheses of early actinopterygian diversification and timescaled phylogenetic tree. (A) A single lineage^16^ or (B) two lineages^19^ cross the extinction boundary, with diversification happening in the earliest Carboniferous. (C) Cryptic diversification happening in the latest Devonian, with many lineages crossing the extinction boundary. (D) Fossil birth-death analysis showing the impact of the inclusion of MCZ VPF 5114 [new taxon name redacted from preprint] and other Famennian/Tournaisian taxa on lineage survivorship over the end-Devonian Mass Extinction. Some clades collapsed for clarity; blue bars associated with nodes indicate 95% highest posterior density for age estimates. Full tree given in Fig. S8. Position of end-Devonian Mass Extinction marked by red vertical line.

Body size represents an important correlate of many aspects of life history and ecology, with many questions centering on impacts of body size during intervals of biotic crisis. Across all marine groups, selectivity on body size is more conspicuous during mass extinctions and their recovery intervals^58^. Decreases of mean body size associated with the Devonian-Carboniferous boundary have previously been reported in many vertebrate clades, including actinopterygians^13^. Our dataset provides a phylogenetically informed assessment of this pattern. While we find a general decrease in body size from the latest Devonian into the early Carboniferous, there is no strong inflection close to the boundary itself (Fig. 4B), and interrogation of size distribution in boundary-crossing lineages suggests that most were mid-sized (Fig. 4A). More densely sampled phylogenetic hypotheses will be necessary for assessing impacts of the Devonian-Carboniferous extinction on the trajectory of body-size evolution in ray-finned fishes.

**Figure 4.**
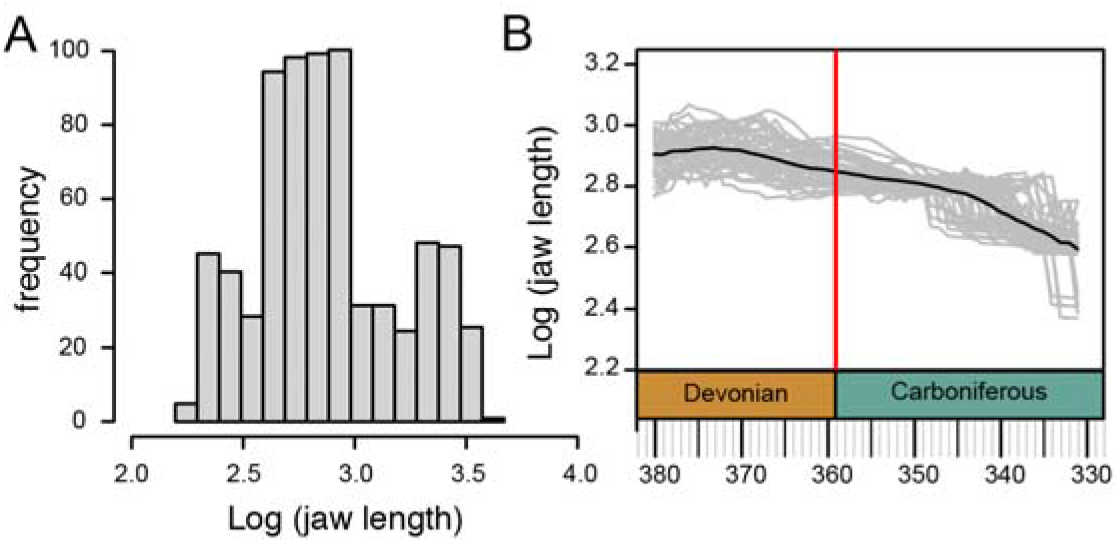
Patterns in early ray-finned fish size around the Devonian-Carboniferous boundary. Size represented by log-transformed lower jaw lengths. (A) Size distribution of boundary-crossing lineages estimated from a sample of 50 trees drawn from the posterior distribution of a Bayesian phylogenetic analysis using the fossilized birth-death prior. (B) Median size estimated at 1 Myr intervals (values inferred along branches plus any coincident tips) during the Late Devonian and early-mid Carboniferous. Black line indicates mean of 50 trajectories (individual grey lines) based on 50 trees drawn from the posterior sample. Position of end-Devonian Mass Extinction marked by red vertical line.

Importantly, the features that nest MCZ VPF 5114 [new taxon name redacted from preprint] and some other Late Devonian taxa within clades composed mostly of younger taxa do not relate to the substantial mandibular and locomotor innovations interpreted as evidence of increased ecological diversity among Carboniferous taxa^9–11,28^. Thus, the initial diversification of actinopterygians appears to show decoupling of cladogenetic events, which took place prior to the end-Devonian extinction, from subsequent ecomorphological and trophic divergence, which took place in the early Carboniferous. Such offsets between lineage divergence and phenotypic diversification are a common feature of vertebrate evolution^59,60^. A similar pattern also seems to characterize the radiation of placental mammals^61,62^. Evidence suggests this might be a common evolutionary trajectory for vertebrate lineages across the Devonian-Carboniferous boundary, with both tetrapods and lungfishes showing substantial lineage diversification before, and morphological divergence after, the extinction^30–35^.

## Methods

### CT scanning

MCZ (Harvard Museum of Comparative Zoology) VPF 5114 was scanned on a Nikon XT 225 ST micro CT scanner at the CTEES facility, Department of Earth and Environmental Sciences, University of Michigan. Details of the scan: voltage = 130 kV; current = 115 μA; exposure = 4 s; frames per projection = 2; projections = 3141; filter = 1 mm Cu. After scanning, data were segmented in Mimics (http://biomedical.materialise.com/mimics; Materialise, Leuven, Belgium). Surface meshes were then exported into and imaged in Blender (http://blender.org; Stitching Blender Foundation, Amsterdam, the Netherlands).

### Phylogenetic analysis

Analyses were performed in TNT v1.5^63^ and MrBayes v.3.2.6^64^ using a dataset modified from Latimer & Giles^50^ and Figueroa et al.^65^ with the addition of ten taxa, giving a total of 121 taxa and 292 characters. Full details are given in the Supplementary Information. An equally weighted parsimony analysis was conducted in TNT using the following settings: hold 200000; xmult=level10; bbreak=fillonly. The outgroup was set as *Dicksonoteus arcticus*, and the following topology constraint was applied: *[Dicksonosteus [Entelognathus [Acanthodes, Cladodoides, Ozarcus*][ingroup]]]. Bremer decay values were calculated in TNT, and an agreement subtree was calculated in PAUP* 4.0a169^66^. We conducted a Bayesian analysis with no stratigraphic information and identical constraints on outgroup taxa, discarding the first third of the run as burnin. To infer the timescale of early actinopterygian diversification, we also conducted an additional Bayesian analysis under the fossil-birth-death (FBD) prior. Taxa were assigned uniform age priors matching the duration of the shortest temporal interval to which they could be assigned (e.g., conodont or ammonoid zones for many Devonian-Carboniferous taxa). In the absence of meaningful prior estimates of key parameters like preservation, extinction, and speciation rates, we adopted uninformative priors suggested by Matzke & Wright^67^. In addition to constraints among non-osteichthyans, we applied a topological constraint for osteichthyan interrelationships matching the strict consensus from parsimony analyses. Analyses were run for a total of 6,000,000 generations, with two chains.

## Data availability

The specimen described in this study is reposited in the collections of the. Harvard Museum of Comparative Zoology. The CT raw projection series, reconstructed .TIFF stack, and .OBJ file of all segmented 3D objects are available on Morphosource.org (https://www.morphosource.org/concern/biological_specimens/000417029). During the review process, the Mimics file can be downloaded upon request from the Department of Vertebrate Paleontology at the Museum of Comparative Zoology; the identities of anyone who downloads the data, including reviewers, will not be made available to the authors of the paper. All other data associated with this paper are included in Supplementary Data.

## Acknowledgments

We thank the Harvard Museum of Comparative Zoology for specimen access and especially Jessica Cundiff for organizing the loan of material that formed the basis of this study. We also thank Emma Bernard, David Gelsthorpe, Zerina Johanson, Lindsay Loughtman, Paul Shephard and Stig Walsh for facilitating access to comparative collections; Graeme Lloyd for helpful discussions; and Christina Byrd for assistance with data archiving. SG was supported by a Royal Society Dorothy Hodgkin Research Fellowship (DH160098). This study includes data produced in the CTEES facility at University of Michigan, supported by the Department of Earth and Environmental Sciences and College of Literature, Science, and the Arts.

## Author Contributions

MF and SP designed research; SG, MF, KF and RW performed research and analysed data; SG and MF wrote the paper and designed figures; all authors edited text.

## Competing Interest Statement

The authors declare no competing interests.

## Notes

### Competing Interest Statement

The authors have declared no competing interest.

### Summary of Updates

We have made some edits to the text, mostly making language more concise and clarifying some points. We have moved a figure to the supplement and rearranged the figure order so that it is more logical, and added a panel to a main text figure to show the contrasting models of early actinopterygian radiation.

