## Supplementary information for "High lineage survivorship across the end-Devonian Mass Extinction suggested by a remarkable new Late Devonian actinopterygian"

^a^School of Geography Earth and Environmental Sciences, University of Birmingham, Birmingham B15 2TT, UK; ^b^Department of Earth Sciences, Natural History Museum, Cromwell Road, London SW7 5BD, UK; ^c^Department of Earth and Environmental Sciences, University of Michigan, 1105 N University Ave, Ann Arbor, MI 48109, USA.; ^d^GeoZentrum Nordbayern, Friedrich-Alexander-Universität Erlangen-Nürnberg, Loewenichstraße 28, 91054 Erlangen Germany; ^e^Museum of Comparative Zoology and Department of Organismic and Evolutionary Biology, Harvard University, 26 Oxford Street, Cambridge, MA 02138, USA; ^f^Museum of Paleontology and Department of Earth and Environmental Sciences, University of Michigan, 1105 N University Ave, Ann Arbor, MI 48109, USA.

Supplementary Information Text

**Phylogenetic analysis**

**General notes.** This character-by-taxon matrix is a composite of Figueroa et al. (2021) and Latimer and Giles (2018), which are both derived from Giles et al. (2017). In addition to the new taxon described here, nine taxa have been added to increase sampling of Givetian (*Stegotrachelus finlayi*: Gardiner, 1963), Famennian (*Cuneognathus gardineri*: Friedman & Blom 2006; *Kentuckia hlavini*: Dunkle 1964; Feldman 1996; *Krasnoyarichthys jesseni*: Prokofiev 2002; *Limnomis delayni*: Daeschler 2000; *Moythomasia perforata*: Gross 1942, Choo 2015; Swartz, 2009) and Tournaisian (*Avonichthys manskyi*: Wilson et al. 2019; *Lambeia pectinatus*: Mickle 2017; *Novogonatodus kasantsevae*: Long 1998; Holland et al. 2007;) taxa. The resulting data matrix has 292 characters and 121 taxa.

**Character list**

**1. [G 1] Large dermal plates**

(Forey, 1980; Gardiner, 1984; Zhu & Schultze, 2001; Zhu et al., 2001; Zhu & Yu, 2002; Zhu et al., 2006; Friedman, 2007; Brazeau, 2009; Zhu et al., 2009; Friedman & Brazeau; 2010; Davis et al., 2012; Zhu et al., 2013; Brazeau & Friedman, 2014; Giles et al., 2015b,c; Giles et al., 2017; Argyriou et al., 2018; Latimer & Giles, 2018; Wilson et al., 2019; Figueroa et al., 2021.)

0 absent

1 present

**2. [G 2] Sensory lines**

(Brazeau, 2009; Zhu et al., 2013; Giles et al., 2015b; Giles et al., 2017; Argyriou et al., 2018; Latimer & Giles, 2018; Wilson et al., 2019; Figueroa et al., 2021. )

0 preserved as open grooves

1 pass through canals

**3. Premaxilla as distinct ossification**

(Hurley et al. 2007; Xu et al. 2014; Giles et al., 2017; Argyriou et al., 2018; Latimer & Giles, 2018; Wilson et al., 2019; Figueroa et al., 2021. Typically, the premaxilla is a short, paired or median bone that contributes to the orbital margin anterior to the maxilla. However, considerable variation is present, and we have attempted to consistently code this variation as laid out here and in the following character descriptions. A premaxilla may be completely absent (e.g. *Acipenser*, *Cyranorhis*) or dorsally expanded into a midline bone (possibly fused with the rostral; e.g. *Bobasatrania*, *Styracopterus*). These two latter states are coded as ?1? here.)

0 present

1 absent

**4. [CH 1; G3] Premaxillae, contact at midline**

(Cloutier & Ahlberg, 1996; Taverne, 1997; Schultze & Cumbaa, 2001; Zhu & Schultze, 2001; Zhu & Yu, 2002; Cloutier & Arratia, 2004; Friedman & Blom, 2006; Zhu et al., 2006; Friedman, 2007; Long et al., 2008; Swartz, 2009; Choo, 2011; Giles et al., 2017; Argyriou et al., 2018; Latimer & Giles, 2018; Wilson et al., 2019; Figueroa et al., 2021. Coded as inapplicable in taxa lacking any ossification in the position typically occupied by the premaxilla (e.g. *Acipenser*, *Cyranorhis*) and where the premaxilla appears fused with the rostral (e.g. *Bobasatrania*, *Styracopterus*).)

0 present

1 absent

**5. Premaxilla fused at midline**

(Xu et al., 2011; Xu et al., 2015; Xu & Zhao, 2016; Giles et al., 2017; Argyriou et al., 2018; Latimer & Giles, 2018; Wilson et al., 2019; Figueroa et al., 2021. Coded as inapplicable in taxa lacking any ossification in the position typically occupied by the premaxilla (e.g. *Acipenser*, *Cyranorhis*) and where the premaxilla appears fused with the rostral (e.g. *Bobasatrania*, *Styracopterus*) and where the premaxilla appears fused with the rostral (e.g. *Bobasatrania*, *Styracopterus*).)

0 absent

1 present

**6. [G 4] Premaxilla**

(Friedman, 2007; Giles et al., 2015b; Giles et al., 2017; Argyriou et al., 2018; Latimer & Giles, 2018; Wilson et al., 2019; Figueroa et al., 2021. Coded as inapplicable in taxa lacking any ossification in the position typically occupied by the premaxilla (e.g. *Acipenser*, *Cyranorhis*) and in taxa where the premaxillae do not contact at the midline.)

0 Reaches or extends past anterior margin of orbit

1 Confined to region anterior to orbit

**7. [G 5] Premaxilla contributes to orbital margin**

(Cloutier & Ahlberg, 1996; Schultze & Cumbaa, 2001; Zhu & Schultze, 2001; Zhu et al., 2001; Zhu & Yu, 2002; Cloutier & Arratia, 2004; Zhu et al., 2006; Friedman, 2007; Long et al., 2008; Swartz, 2009; Zhu et al., 2009; Xu & Gao, 2011; Zhu et al., 2013; Xu et al., 2014; Giles et al., 2017; Argyriou et al., 2018; Latimer & Giles, 2018; Wilson et al., 2019; Figueroa et al., 2021. Coded as inapplicable in taxa lacking any ossification in the position typically occupied by the premaxilla (e.g. *Acipenser*, *Cyranorhis*), where the premaxilla appears fused with the rostral (e.g. *Bobosatrania*, *Styracopterus*), and where the premaxilla is restricted anterior to the orbit.)

0 absent

1 present

**8. Teeth on premaxillae**

(Cloutier & Arratia 2004, Xu et al. 2014; Giles et al., 2017; Argyriou et al., 2018; Latimer & Giles, 2018; Wilson et al., 2019; Figueroa et al., 2021. Coded as inapplicable in taxa lacking any ossification in the position typically occupied by the premaxilla (e.g. *Acipenser*, *Cyranorhis*).)

0 present

1 absent

**9. Mobile premaxilla**

(Arratia 1999; Cavin & Suteethorn 2006; Hurley et al. 2007; Giles et al., 2017; Argyriou et al., 2018; Latimer & Giles, 2018; Wilson et al., 2019; Figueroa et al., 2021. Coded as inapplicable in taxa lacking any ossification in the position typically occupied by the premaxilla (e.g. *Acipenser*, *Cyranorhis*) and where the premaxilla appears fused with the rostral (e.g. *Bobasatrania*, *Styracopterus*))

0 absent

1 present

**10. Olfactory nerve pierces premaxilla**

(Grande 2010; Xu et al., 2015; Xu & Shen, 2015; Xu & Zhao, 2016; Giles et al., 2017; Argyriou et al., 2018; Latimer & Giles, 2018; Wilson et al., 2019; Figueroa et al., 2021. Coded as inapplicable in taxa lacking any ossification in the position typically occupied by the premaxilla (e.g. *Acipenser*, *Cyranorhis*) and where the premaxilla appears fused with the rostral (e.g. *Bobasatrania*, *Styracopterus*).)

0 absent

1 present

**11. Nasal process of premaxilla**

(Gardiner & Schaeffer 1989; Olsen & McCune 1991; Gardiner et al. 1996; Gardiner et al. 2005; Cavin & Suteethorn 2006; Hurley et al. 2007; Grande 2010; LÓpez-Arbarello 2012; Xu & Wu 2012; Xu et al. 2014; Bermúdez-Rochas & Poyato-Ariza 2015; Poyato-Ariza 2015; Thies & Waschkewitz, 2015; Xu & Shen, 2015; Gibson 2016; Xu & Zhao, 2016; Giles et al., 2017; López-Arbarello & Sferco 2018; Argyriou et al., 2018; Latimer & Giles, 2018; Wilson et al., 2019; Figueroa et al., 2021. Coded as inapplicable in taxa lacking any ossification in the position typically occupied by the premaxilla (e.g. *Acipenser*, *Cyranorhis*) and where the premaxilla appears fused with the rostral (e.g. *Bobasatrania*, *Styracopterus*).)

0 absent

1 short

2 long, reaches skull roof

**12. Sensory canal on premaxilla**

(Giles et al., 2017; Argyriou et al., 2018; Latimer & Giles, 2018; Wilson et al., 2019; Figueroa et al., 2021. Coded as inapplicable in taxa lacking any ossification in the position typically occupied by the premaxilla (e.g. *Acipenser*, *Cyranorhis*) and where the premaxilla appears fused with the rostral (e.g. *Bobasatrania*, *Styracopterus*).)

0 present

1 absent

**13. [CH 3; G 6] Postrostrals (element[s] immediately anterior to frontals but not in contact with premaxillae)**

(Cloutier & Ahlberg, 1996; Taverne, 1997; Lund, 2000; Schultze & Cumbaa, 2001; Zhu & Schultze, 2001; Lund & Poplin, 2002; Cloutier & Arratia, 2004; Friedman & Blom, 2006; Long et al., 2008; Swartz, 2009; Choo, 2011; Xu et al., 2014; Giles et al., 2015b; Giles et al., 2017; Argyriou et al., 2018; Latimer & Giles, 2018; Wilson et al., 2019; Figueroa et al., 2021.)

0 present

1 absent

**14. [CH 4; G 7] Single median dermal bone capping snout**

(Gardiner & Schaeffer, 1989; Taverne, 1997; Friedman & Blom, 2006; Long et al., 2008; Swartz, 2009; Choo, 2011; Giles et al., 2015b; Poyato-Ariza 2015; Giles et al., 2017; Argyriou et al., 2018; Latimer & Giles, 2018; Wilson et al., 2019; Figueroa et al., 2021.)

0 absent

1 present

**15. Median rostal**

(Gardiner et al. 1996; Hurley et al. 2007; Giles et al., 2017; Argyriou et al., 2018; Latimer & Giles, 2018; Wilson et al., 2019; Figueroa et al., 2021.)

0 plate-like

1 tube-like

**16. [G 8] Pores for rostral organ**

(Friedman, 2007; Giles et al., 2015b; Giles et al., 2017; Argyriou et al., 2018; Latimer & Giles, 2018; Wilson et al., 2019; Figueroa et al., 2021.)

0 absent

1 present

**17. [CH 8; G 10] Nasal bone as single consolidated ossification (i.e. bone(s) carrying supraorbital canal between premaxilla and anterior margin of frontals)**

(Taverne, 1997; Schultze & Cumbaa, 2001; Friedman & Blom, 2006; Long et al., 2008; Swartz, 2009; Choo, 2011; Giles et al., 2015b; Giles et al., 2017; Argyriou et al., 2018; Latimer & Giles, 2018; Wilson et al., 2019; Figueroa et al., 2021.)

0 absent

1 present

**18. Contact of nasals on midline**

(Giles et al., 2017; Argyriou et al., 2018; Latimer & Giles, 2018; Wilson et al., 2019; Figueroa et al., 2021.)

0 separated by dermal bones

1 contacting or separated by gap unfilled by bone

**19. Nasal contributes to orbital margin**

(Xu & Wu 2012; Xu et al. 2014; Xu & Zhao, 2016; Giles et al., 2017; López-Arbarello & Sferco 2018; Argyriou et al., 2018; Latimer & Giles, 2018; Wilson et al., 2019; Figueroa et al., 2021.)

0 absent

1 present

**20. [CH 57; G 11] Mesial margin of (anterior) nasal**

(Lund et al., 1995; Ahlberg & Johanson, 1998; Ahlberg et al., 2000; Lund, 2000; Poplin & Lund, 2000; Schultze & Cumbaa, 2001; Lund & Poplin, 2002; Cloutier & Arratia, 2004; Zhu & Ahlberg, 2004; Daeschler et al., 2006; Long et al., 2006; Zhu et al., 2006; Zhu et al., 2009; Choo, 2011; Giles et al., 2015b; Giles et al., 2017; López-Arbarello & Sferco 2018; Argyriou et al., 2018; Latimer & Giles, 2018; Wilson et al., 2019; Figueroa et al., 2021.)

0 not notched

1 notched

**21. [CH 6; G12] Posterior nostril in complete communication with orbital fenestra**

(Friedman & Blom, 2006; Long et al., 2008; Choo, 2011; Giles et al., 2015b; Giles et al., 2017; Argyriou et al., 2018; Latimer & Giles, 2018; Wilson et al., 2019; Figueroa et al., 2021. This character refers to the opening of the posterior nostril with reference to the surrounding dermal bones.)

0 absent

1 present

**22. [CH 7; G 13] Posterior nostril: contribution to margin by premaxillae**

(Friedman & Blom, 2006; Long et al., 2008; Choo, 2011; Giles et al., 2015b; Giles et al., 2017; Argyriou et al., 2018; Latimer & Giles, 2018; Wilson et al., 2019; Figueroa et al., 2021.)

0 absent

1 present

**23. [G 14] Tectals (sensu Cloutier & Ahlberg 1996, not counting the posterior tectal of Jarvik)** (Lund et al., 1995; Cloutier & Ahlberg, 1996; Lund, 2000; Schultze & Cumbaa, 2001; Zhu & Schultze, 2001; Zhu et al., 2001; Lund & Poplin, 2002; Zhu & Yu, 2002; Cloutier & Arratia, 2004; Zhu et al., 2006; Friedman, 2007; Swartz, 2009; Zhu et al., 2009; Zhu et al., 2013; Giles et al., 2015b; Giles et al., 2017; Argyriou et al., 2018; Latimer & Giles, 2018; Wilson et al., 2019; Figueroa et al., 2021. )

0 absent

1 present

**24. [CH 75; G 36] Dermal intracranial joint**

(Cloutier & Ahlberg, 1996; Ahlberg & Johanson, 1998; Zhu & Ahlberg, 2004; Zhu & Schultze, 2001; Zhu et al., 2001; Zhu & Yu, 2002; Daeschler et al., 2006; Long et al., 2006; Zhu et al., 2006; Friedman, 2007; Brazeau, 2009; Zhu et al., 2009; Choo, 2011; Davis et al., 2012; Zhu et al., 2013; Giles et al., 2015b; Giles et al., 2017; Argyriou et al., 2018; Latimer & Giles, 2018; Wilson et al., 2019; Figueroa et al., 2021.)

0 absent

1 present

**25. [CH 9; G 15] Pineal foramen**

(Cloutier & Ahlberg, 1996; Taverne, 1997; Schultze & Cumbaa, 2001; Zhu & Schultze, 2001; Zhu & Yu, 2002; Friedman & Blom, 2006; Friedman, 2007; Long et al., 2008; Brazeau, 2009; Swartz, 2009; Davis et al., 2012; Zhu et al., 2013; Xu et al., 2014; Giles et al., 2015b,c; Giles et al., 2017; Argyriou et al., 2018; Latimer & Giles, 2018; Wilson et al., 2019; Figueroa et al., 2021. A pineal foramen is variably present in *Cheirolepis canadensis* (Pearson & Westoll, 1979; Arratia & Cloutier, 1996), *C. trailli* (Pearson & Westoll, 1979), *Kentuckia deani* (Rayner, 1951) and *Meemannia* (Zhu et al., 2010), and these taxa are coded '0/1' to reflect this polymorphism.)

0 present

1 absent

**26. [G 16] Pineal eminence**

(Friedman, 2007; Zhu et al., 2009; Giles et al., 2015b; Giles et al., 2017; Argyriou et al., 2018; Latimer & Giles, 2018; Wilson et al., 2019; Figueroa et al., 2021. Can only be coded in taxa that lack a pineal foramen. )

0 absent

1 present

**27. [CH 10; G17] Shape of parietals (sarcopterygian postparietals):**

(Dietze, 2000; Schultze & Cumbaa, 2001; Cloutier & Arratia, 2004; Friedman & Blom, 2006; Long et al., 2008; Swartz, 2009; Choo, 2011; Xu et al., 2014; Giles et al., 2015b; Giles et al., 2017; Argyriou et al., 2018; Latimer & Giles, 2018; Wilson et al., 2019; Figueroa et al., 2021.)

0 rectangular, with long axis parallel to midline

1 quadrate

**28. [CH 11; G 18] Relative lengths of frontals and parietals (sarcopterygian parietals and postparietals)**

(Lund et al., 1995; Taverne, 1997; Dietze, 2000; Lund, 2000; Poplin & Lund, 2000; Schultze & Cumbaa, 2001; Lund & Poplin, 2002; Cloutier & Arratia, 2004; Friedman & Blom, 2006; Zhu et al., 2006; Long et al., 2008; Swartz, 2009; Choo, 2011; López-Arbarello 2012; Xu et al., 2014; Bermúdez-Rochas & Poyato-Ariza 2015; Thies & Waschkewitz, 2015; Gibson 2016; Giles et al., 2015b; Giles et al., 2017; López-Arbarello & Sferco 2018; Argyriou et al., 2018; Latimer & Giles, 2018; Wilson et al., 2019; Figueroa et al., 2021.)

0 frontal shorter than parietal

1 frontal approximately equal to parietal

2 frontal longer than parietal

**29. Frontals broad posteriorly and tapering anteriorly**

(Arratia 1999; López-Arbarello 2012; Bermúdez-Rochas & Poyato-Ariza 2015; Thies & Waschkewitz, 2015; Gibson 2016; Giles et al., 2017; López-Arbarello & Sferco 2018; Argyriou et al., 2018; Latimer & Giles, 2018; Wilson et al., 2019; Figueroa et al., 2021.)

0 absent

1 present

**30. [G 19] Anterior pit line**

(Giles et al., 2015b,c; Giles et al., 2017; Argyriou et al., 2018; Latimer & Giles, 2018; Wilson et al., 2019; Figueroa et al., 2021. Although not figured, an anterior pit line is described for *Miguashaia* (Cloutier 1996).)

0 absent

1 present

**31. [G 20] Otic canal extends through parietals**

(Giles et al., 2015b,c; Giles et al., 2017; Argyriou et al., 2018; Latimer & Giles, 2018; Wilson et al., 2019; Figueroa et al., 2021.)

0 absent

1 present

**32. Junction between supraorbital and infraorbital canals**

(López-Arbarello 2012; Bermúdez-Rochas & Poyato-Ariza 2015; Poyato-Ariza 2015; Giles et al., 2017; López-Arbarello & Sferco 2018; Argyriou et al., 2018; Latimer & Giles, 2018; Wilson et al., 2019; Figueroa et al., 2021. The supraorbital canal may terminate in the frontal/parietal, or it may become confluent with the infraorbital canal. The exact position of this junction is highly variable, and typically occurs in the region of the frontal, dermosphenotic, dermopterotic.)

0 absent

1 present

**33. Anterior branch of infraorbital sensory canal**

(Giles et al., 2017; Argyriou et al., 2018; Latimer & Giles, 2018; Wilson et al., 2019; Figueroa et al., 2021. In some taxa (e.g. *Dapedium*), the infraorbital canal continues anteriorly above the orbit a short way.)

0 absent

1 present

**34. [G 21] Tabular**

(Lund et al., 1995; Cloutier & Ahlberg, 1996; Schultze & Cumbaa, 2001; Zhu & Schultze, 2001; Cloutier & Arratia, 2004; Long et al., 2008; Swartz, 2009; Giles et al., 2015b; Giles et al., 2017; Argyriou et al., 2018; Latimer & Giles, 2018; Wilson et al., 2019; Figueroa et al., 2021.)

0 present

1 absent

**35. [G 22] Tabular pit line**

(Giles et al., 2015b,c; Giles et al., 2017; Argyriou et al., 2018; Latimer & Giles, 2018; Wilson et al., 2019; Figueroa et al., 2021.)

0 absent

1 present

**36. [CH 64; G 28] Number of bones carrying otic portion of lateral line canal between dermosphenotic and posterior edge of skull roof.**

(Gardiner & Schaeffer 1989; Cloutier & Arratia, 2004; Hurley et al. 2007; Choo, 2011; Giles et al., 2015b; Xu & Zhao, 2016; Giles et al., 2017; Argyriou et al., 2018; Latimer & Giles, 2018; Wilson et al., 2019; Figueroa et al., 2021. This character is reformulated from Choo's character 'Dermopterotic: present/absent'. Rather than designating bones as an intertemporal (or supratemporal or dermosphenotic) a priori, we consider the number of bones carrying the otic portion of the lateral line canal between the dermosphenotic and the posterior edge of the skull roof. Where two bones are present, these are treated as the intertemporal and supratemporal; where only one is present, this is treated as the dermopterotic. Anamestic bones between the dermosphenotic and frontal are not included in this count.) )

0 at least two (i.e. intertemporal and supratemporal)

1 one (i.e. dermopterotic)

**37. [CH 12; G 23] Intertemporal: relative length**

(Taverne, 1997; Friedman & Blom, 2006; Choo, 2011; Giles et al., 2015b; Giles et al., 2017; Argyriou et al., 2018; Latimer & Giles, 2018; Wilson et al., 2019; Figueroa et al., 2021. Coded as inapplicable in taxa with a dermopterotic. Coded as inapplicable in taxa with a dermopterotic.)

0 shorter than supratemporal

1 of similar length to supratemporal

2 longer than supratemporal

**38. [CH 13; G 24] Intertemporal: contact with supratemporal anterior to that between frontal and parietal**

(Friedman & Blom, 2006; Choo, 2011; Giles et al., 2015b; Giles et al., 2017; Argyriou et al., 2018; Latimer & Giles, 2018; Wilson et al., 2019; Figueroa et al., 2021. Coded as inapplicable in taxa with a dermopterotic.)

0 absent

1 present

**39. [G 27] Intertemporal contacts nasal**

(Xu & Gao, 2011; Xu et al., 2014; Giles et al., 2015b; Giles et al., 2017; Argyriou et al., 2018; Latimer & Giles, 2018; Wilson et al., 2019; Figueroa et al., 2021. Coded as inapplicable in taxa with a dermopterotic.)

0 absent

1 present

**40. [CH 69; G 29] Supratemporal: narrow anterolateral flange forming ventral margin of spiracular opening**

(Choo, 2011; Giles et al., 2015b; Giles et al., 2017; Argyriou et al., 2018; Latimer & Giles, 2018; Wilson et al., 2019; Figueroa et al., 2021. Coded as inapplicable in taxa with a dermopterotic.)

0 absent

1 present

**41. Parietal fused to dermopterotic**

(Xu & Gao 2011; Xu et al. 2014; Giles et al., 2017; Argyriou et al., 2018; Latimer & Giles, 2018; Wilson et al., 2019; Figueroa et al., 2021. Coded as inapplicable in taxa with a separate intertemporal and supertemporal, and in taxa lacking these bones entirely.)

0 absent

1 present

**42. Bone carrying otic portion of lateral line canal extends past posterior margin of parietals**

(Lu et al. 2016; Giles et al., 2017; Argyriou et al., 2018; Latimer & Giles, 2018; Wilson et al., 2019; Figueroa et al., 2021.)

0 absent

1 present

**43. [CH 15; G 30] Number of paired extrascapulars**

(Gardiner & Schaeffer, 1989; Lund et al., 1995; Cloutier & Ahlberg, 1996; Grande & Bemis 1998; Coates, 1998; Lund, 2000; Poplin & Lund, 2000; Schultze & Cumbaa, 2001; Zhu & Schultze, 2001; Lund & Poplin, 2002; Cloutier & Arratia, 2004; Friedman & Blom, 2006; Long et al., 2008; Swartz, 2009; Choo, 2011; López-Arbarello 2012; Zhu et al., 2013; Bermúdez-Rochas & Poyato-Ariza 2015; Poyato-Ariza 2015; Giles et al., 2015b; Thies & Waschkewitz, 2015; Xu et al., 2015; Gibson 2016; Xu & Zhao, 2016; Giles et al., 2017; López-Arbarello & Sferco 2018; Argyriou et al., 2018; Latimer & Giles, 2018; Wilson et al., 2019; Figueroa et al., 2021. )

0 one pair

1 two pairs

2 three or more pairs

**44. [G 31] Extrascapular reaches lateral edge of skull roof**

(Giles et al., 2015b,c; Giles et al., 2017; Argyriou et al., 2018; Latimer & Giles, 2018; Wilson et al., 2019; Figueroa et al., 2021. The skull roof of *Moythomasia* as shown in Fig. 103 (Gardiner, 1984) is a restoration (see also Choo 2015: fig. 13). The only skull roof directly figured by Gardiner does not preserve the extrascapulars in situ (fig. 83). However, in specimens viewed by us, as well as in published photos of articulated material (e.g. Choo 2015: fig 8), the lateral extrascapular of *M. durgaringa* is clearly excluded from the lateral margin of the skull roof.)

0 absent

1 present

**45. [CH 71; G 32] Single median extrascapular**

(Dietze, 2000; Cloutier & Arratia, 2004; Long et al., 2008; Swartz, 2009; Choo, 2011; Xu & Gao, 2011; Zhu et al., 2013; Xu et al., 2014; Giles et al., 2015b; Giles et al., 2017; Argyriou et al., 2018; Latimer & Giles, 2018; Wilson et al., 2019; Figueroa et al., 2021.)

0 present

1 absent

**46. [G 33] Extrascapulae contact each other at midline**

(Giles et al., 2015b,c; Giles et al., 2017; Argyriou et al., 2018; Latimer & Giles, 2018; Wilson et al., 2019; Figueroa et al., 2021. Inapplicable for taxa that possess a median extrascapular, as it is logically impossible for the lateral extrascapulae to meet in the midline.)

0 absent

1 present

**47. [CH 70; G 34] Medially-directed branch of sensory canal in extrascapulae**

(Choo, 2011; Giles et al., 2015b; Giles et al., 2017; Argyriou et al., 2018; Latimer & Giles, 2018; Wilson et al., 2019; Figueroa et al., 2021.)

0 present

1 absent

**48. [G 35] Extratemporal**

(Cloutier & Ahlberg, 1996; Ahlberg & Johanson, 1998; Zhu & Schultze, 2001; Zhu et al., 2001; Zhu & Yu, 2002; Zhu & Ahlberg, 2004; Daeschler et al., 2006; Long et al., 2006; Zhu et al., 2006; Friedman, 2007; Zhu et al., 2009; Giles et al., 2015b; Giles et al., 2017; Argyriou et al., 2018; Latimer & Giles, 2018; Wilson et al., 2019; Figueroa et al., 2021.)

0 absent

1 present

**49. [CH 59; G 38] Antorbital bone**

(Gardiner & Schaeffer 1989; Lund et al. 2000; Cloutier & Arratia, 2004; Hurley et al. 2007; Choo, 2011; Xu & Gao, 2011; Xu et al., 2014; Giles et al., 2015b; Giles et al., 2017; López-Arbarello & Sferco 2018; Argyriou et al., 2018; Latimer & Giles, 2018; Wilson et al., 2019; Figueroa et al., 2021.)

0 absent

1 present

**50. Tube-like canal bearing anterior arm of antorbital:**

(Grande 2010; Xu & Wu 2012; Xu et al. 2014; Poyato-Ariza 2015; Xu & Shen, 2015; Xu & Zhao, 2016; Giles et al., 2017; López-Arbarello & Sferco 2018; Argyriou et al., 2018; Latimer & Giles, 2018; Wilson et al., 2019; Figueroa et al., 2021.)

0 absent

1 present

**51. [CH 61; G 39] Infraorbitals**

(Cloutier & Arratia, 2004; Gardiner et al., 2005; Choo, 2011; Xu & Gao, 2011; Xu et al., 2014; Giles et al., 2015b; Giles et al., 2017; López-Arbarello & Sferco 2018; Argyriou et al., 2018; Latimer & Giles, 2018; Wilson et al., 2019; Figueroa et al., 2021.)

0 one

1 two

2 more than two

**52. [CH 16; G 40] Anterior expansion of lacrimal**

(Taverne, 1997; Friedman & Blom, 2006; Long et al., 2008; Swartz, 2009; Choo, 2011; Giles et al., 2015b; Giles et al., 2017; Argyriou et al., 2018; Latimer & Giles, 2018; Wilson et al., 2019; Figueroa et al., 2021.)

0 absent

1 present

**53. [CH 17; G 41] Notch in anterior margin of jugal**

(Cloutier & Arratia, 2004; Friedman & Blom, 2006; Long et al., 2008; Swartz, 2009; Choo, 2011; Giles et al., 2015b; Giles et al., 2017; Xu et al., 2014; Argyriou et al., 2018; Latimer & Giles, 2018; Wilson et al., 2019; Figueroa et al., 2021. )

0 absent

1 present

**54. [CH 18; G 42] Suborbitals (non-canal bearing ossifications separating jugal and maxilla)** (Gardiner & Schaeffer 1989; Olsen & McCune 1991; Taverne, 1997; Schultze & Cumbaa, 2001; Friedman & Blom, 2006; Long et al., 2008; Choo, 2011; López-Arbarello 2012; Xu & Gao 2011; Bermúdez-Rochas & Poyato-Ariza 2015; Poyato-Ariza 2015; Giles et al., 2015b; Thies & Waschkewitz, 2015; Xu et al., 2015; Gibson 2016; Xu & Zhao, 2016; Giles et al., 2017; López-Arbarello & Sferco 2018; Argyriou et al., 2018; Latimer & Giles, 2018; Wilson et al., 2019; Figueroa et al., 2021.)

0 absent

1 one

2 two

3 three or more

**55. [G 43] Multiple rami of infraorbital canal in jugal**

(Giles et al., 2015b,c; Giles et al., 2017; Argyriou et al., 2018; Latimer & Giles, 2018; Wilson et al., 2019; Figueroa et al., 2021. Multiple branches radiate from the infraorbital canal in the jugal of many Carboniferous and younger actinopterygians, as well as the new taxon described here.)

0 absent

1 present

**56. [CH 54; G 25] Dermosphenotic with distinct posterior ramus**

(Gardiner & Schaeffer, 1989; Coates, 1998; Schultze & Cumbaa, 2001; Cloutier & Arratia, 2004; Friedman & Blom, 2006; Zhu et al., 2006; Long et al., 2008; Zhu et al., 2009; Choo, 2011; Giles et al., 2015b; Xu et al., 2015; Giles et al., 2017; Argyriou et al., 2018; Latimer & Giles, 2018; Wilson et al., 2019; Figueroa et al., 2021. The dermosphenotic of *Moythomasia* illustrated by (Gardiner 1984, fig 69) lacks a posterior limb, but this is from a small individual and most likely reflects ontogenetic variability, with a posterior limb being developed in larger individuals (B. Choo, pers. comm.; Choo 2015: fig. 8). The posterior limb of the dermosphenotic is variably developed in *Mesopoma* (Coates, 1999), so this taxon is scored '0/1' to reflect this polymorphism.)

0 absent

1 present

**57. [CH 14; G 26] Dermosphenotic: contact with frontals blocked by intertemporal or dermopterotic**

(Friedman & Blom, 2006; Choo, 2011; Giles et al., 2015b; Giles et al., 2017; Argyriou et al., 2018; Latimer & Giles, 2018; Wilson et al., 2019; Figueroa et al., 2021.)

0 absent

1 present

**58. Supraorbital**

(Gardiner & Schaeffer 1989; Grande & Bemis 1998; Hurley et al. 2007; Xu & Gao 2011; Xu et al. 2014; Poyato-Ariza 2015; Xu et al., 2015; Xu & Zhao, 2016; Giles et al., 2017; López-Arbarello & Sferco 2018; Argyriou et al., 2018; Latimer & Giles, 2018; Wilson et al., 2019; Figueroa et al., 2021.)

0 absent

1 one or two

2 three or more

**59. Anterior-most infraorbital anterior to orbit (i.e. does not contribute to orbital margin)**

(Olsen & McCune 1991; Cavin & Suteethorn 2006; López-Arbarello 2012; Bermúdez-Rochas & Poyato-Ariza 2015; Thies & Waschkewitz, 2015; Gibson 2016; Giles et al., 2017; López-Arbarello & Sferco 2018; Argyriou et al., 2018; Latimer & Giles, 2018; Wilson et al., 2019; Figueroa et al., 2021.)

0 absent

1 present

**60. Three or more lachrymals**

(Grande 2010; Xu & Wu 2012; Xu et al. 2014, 2015; Bermódez-Rochas & Poyato-Ariza 2015; Poyato-Ariza 2015; Thies & Waschkewitz, 2015; Gibson 2016; Xu & Zhao, 2016; Giles et al., 2017; Argyriou et al., 2018; Latimer & Giles, 2018; Wilson et al., 2019; Figueroa et al., 2021. The first lachrymal is regarded here as the anteriormost canal-bearing bone that contributes to orbital margin.)

0 absent

1 present

**61. Circumorbital ring**

(Wiley 1976; Lopez-Arbarello 2010.Wiley 1976; López-Arbarello 2012; Bermúdez-Rochas & Poyato-Ariza 2015; Poyato-Ariza 2015; Thies & Waschkewitz, 2015; Gibson 2016; Giles et al., 2017; López-Arbarello & Sferco 2018; Argyriou et al., 2018; Latimer & Giles, 2018; Wilson et al., 2019; Figueroa et al., 2021.)

0 Supraorbitals do not contact infraorbitals at the anterior rim of the orbit.

1 Supraorbitals contact infraorbitals, closing the orbit.

**62. [CH 62; G 44] Jugal canal**

(Patterson, 1982; Lauder & Liem, 1983; Gardiner, 1984; Cloutier & Arratia, 2004; Brazeau, 2009; Friedman & Brazeau, 2010; Choo, 2011; Davis et al., 2012; Zhu et al., 2013; Giles et al., 2015b,c; Giles et al., 2017; Argyriou et al., 2018; Latimer & Giles, 2018; Wilson et al., 2019; Figueroa et al., 2021.)

0 absent

1 present

**63. [CH 53; G 45] Dermohyal**

(Patterson, 1982; Gardiner & Schaeffer, 1989; Lund et al., 1995; Cloutier & Ahlberg, 1996; Coates, 1998; Dietze, 2000; Lund, 2000; Schultze & Cumbaa, 2001; Zhu & Schultze, 2001; Zhu et al., 2001; Lund & Poplin, 2002; Zhu & Yu, 2002; Cloutier & Arratia, 2004; Gardiner et al., 2005; Friedman & Blom, 2006; Zhu et al., 2006; Friedman, 2007; Long et al., 2008; Swartz, 2009; Zhu et al., 2009; Choo, 2011; Xu & Gao, 2011; Xu et al., 2014; 2015; Giles et al., 2015b; Xu & Zhao, 2016; Giles et al., 2017; Argyriou et al., 2018; Latimer & Giles, 2018; Wilson et al., 2019; Figueroa et al., 2021. This region of the cheek is missing in *Coccocephalichthys* (Poplin, 1974; Poplin & Véran, 1996), and was presumably removed by Watson (1925) when he first described the specimen. It is unclear from the surviving cast whether a dermohyal and/or accessory operculum were present, and as such this taxon is coded as ???.)

0 absent

1 present

**64. [G 46] Head of dermohyal projects above dorsal margin of operculum**

(Giles et al., 2015b,c; Giles et al., 2017; Argyriou et al., 2018; Latimer & Giles, 2018; Wilson et al., 2019; Figueroa et al., 2021. The dermohyal is not preserved in *Melanecta* (Coates, 1998), but it is clear from the surrounding bones that it would not have projected above the dorsal surface of the operculum.)

0 absent

1 present

**65. [G 47] Dermohyal**

(Gardiner et al., 2005; Coates, 1999; Xu & Gao, 2011; Xu et al., 2014; Giles et al., 2015b; Giles et al., 2017; Argyriou et al., 2018; Latimer & Giles, 2018; Wilson et al., 2019; Figueroa et al., 2021.)

0 fused to hyomandibular

1 separate from hyomandibular

**66. [G 49] Complete enclosure of spiracle by bones bearing otic and infraorbital canals** (Friedman, 2007; Zhu et al., 2009; Giles et al., 2015b; Giles et al., 2017; Argyriou et al., 2018; Latimer & Giles, 2018; Wilson et al., 2019; Figueroa et al., 2021.)

0 absent

1 present

**67. [G 57] Maxilla**

(Zhu & Yu, 2002; Friedman, 2007; Xu et al., 2014, 2015; Giles et al., 2015b; Xu & Zhao, 2016; Giles et al., 2017; López-Arbarello & Sferco 2018; Argyriou et al., 2018; Latimer & Giles, 2018; Wilson et al., 2019; Figueroa et al., 2021.)

0 absent

1 present

**68. [G 58] Expanded dorsal lamina of maxilla**

(Lund et al., 1995; Lund, 2000; Poplin & Lund, 2000; Schultze & Cumbaa, 2001; Zhu & Schultze, 2001; Zhu et al., 2001; Zhu & Yu, 2002; Lund & Poplin, 2002; Cloutier & Arratia, 2004; Zhu et al., 2006; Friedman, 2007; Zhu et al., 2009; Zhu et al., 2013; Giles et al., 2015b,c; Giles et al., 2017; López-Arbarello & Sferco 2018; Argyriou et al., 2018; Latimer & Giles, 2018; Wilson et al., 2019; Figueroa et al., 2021. )

0 absent

1 present

**69. [G 60] Contribution by maxilla to posterior margin of cheek**

(Friedman, 2007; Zhu et al., 2009; Zhu et al., 2013; Giles et al., 2015b,c; Giles et al., 2017; López-Arbarello & Sferco 2018; Argyriou et al., 2018; Latimer & Giles, 2018; Wilson et al., 2019; Figueroa et al., 2021.)

0 absent

1 present

**70. [G 61] Sensory canal/pit line associated with maxilla**

(Grande & Bemis 1998; Friedman, 2007; Zhu et al., 2009; Zhu et al., 2013; Giles et al., 2015b; Giles et al., 2017; Argyriou et al., 2018; Latimer & Giles, 2018; Wilson et al., 2019; Figueroa et al., 2021.)

0 absent

1 present

**71. Teeth on maxilla**

(Poyato-Ariza & Wenz, 2002; Cloutier & Arratia, 2004; López-Arbarello 2012; Xu et al. 2014, 2015; Bermúdez-Rochas & Poyato-Ariza 2015; Poyato-Ariza 2015; Thies & Waschkewitz, 2015; Gibson 2016; Xu & Zhao, 2016; Giles et al., 2017; Cawley & Kriwet 2018; López-Arbarello & Sferco 2018; Argyriou et al., 2018; Latimer & Giles, 2018; Wilson et al., 2019; Figueroa et al., 2021.)

0 present

1 absent

**72. Mobile maxilla in cheek**

(Patterson 1973; Gardiner & Schaeffer 1989; Olsen & McCune 1991; Gardiner et al. 1996; Gardiner et al. 2005; Coates 1999; Hurley et al. 2007; Xu & Gao 2011; Poyato-Ariza 2015; Xu et al. 2014, 2015; Xu & Zhao, 2016; Giles et al., 2017; Argyriou et al., 2018; Latimer & Giles, 2018; Wilson et al., 2019; Figueroa et al., 2021.)

0 absent

1 present

**73. Peg-like anterior process of maxilla**

(Grande 2010; Xu & Wu 2012; Xu et al. 2014; Poyato-Ariza 2015; Giles et al., 2017; Argyriou et al., 2018; Latimer & Giles, 2018; Wilson et al., 2019; Figueroa et al., 2021.)

0 absent

1 present

**74. Posterior maxillary notch**

(Grande & Bemis 1998; Xu & Wu 2012; Xu et al. 2014, 2015. Arratia 2013; Xu & Zhao, 2016; Giles et al., 2017; López-Arbarello & Sferco 2018; Argyriou et al., 2018; Latimer & Giles, 2018; Wilson et al., 2019; Figueroa et al., 2021.)

0 absent

1 present

**75. Supramaxilla**

(Grande & Bemis 1998; Gardiner & Schaeffer 1989; Olsen & McCune 1991; Gardiner et al. 1996; Gardiner et al. 2005; Coates 1999; Hurley et al. 2007; Xu & Gao 2011; Xu et al. 2014, 2015; Bermúdez-Rochas & Poyato-Ariza 2015; Poyato-Ariza 2015; Xu & Shen, 2015; Thies & Waschkewitz, 2015; Gibson 2016; Xu & Zhao, 2016; Giles et al., 2017; López-Arbarello & Sferco 2018; Argyriou et al., 2018; Latimer & Giles, 2018; Wilson et al., 2019; Figueroa et al., 2021.)

0 absent

1 one

2 two

**76. [CH 21; G 63] Course of mandibular canal**

(Friedman & Blom, 2006; Long et al., 2008; Swartz, 2009; Choo, 2011; Giles et al., 2015b; Giles et al., 2017; Argyriou et al., 2018; Latimer & Giles, 2018; Wilson et al., 2019; Figueroa et al., 2021.)

0 traces ventral margin of jaw along entire length

1 arches dorsally in anterior half of jaw

**77. [G 64] Mandibular canal reaches anterior margin of mandible**

(Giles et al., 2015b,c; Giles et al., 2017; Argyriou et al., 2018; Latimer & Giles, 2018; Wilson et al., 2019; Figueroa et al., 2021.)

0 present

1 absent

**78. [CH 74; G 65] Mandibular canal**

(Patterson, 1982; Cloutier & Ahlberg, 1996; Coates, 1998; Schultze & Cumbaa, 2001; Zhu & Schultze, 2001; Zhu et al., 2001; Zhu & Yu, 2002; Cloutier & Arratia, 2004; Zhu et al., 2006; Friedman, 2007; Zhu et al., 2009; Choo, 2011; Zhu et al., 2013; Giles et al., 2015b; Giles et al., 2017; Argyriou et al., 2018; Latimer & Giles, 2018; Wilson et al., 2019; Figueroa et al., 2021.)

0 primarily carried by infradentaries

1 primarily carried by dentary

**79. [G 66] Relative length of dentary**

(Ahlberg & Johanson, 1998; Zhu et al., 2001; Zhu & Yu, 2002; Zhu & Ahlberg, 2004; Friedman, 2007; Zhu et al., 2009; Giles et al., 2015b; Giles et al., 2017; Argyriou et al., 2018; Latimer & Giles, 2018; Wilson et al., 2019; Figueroa et al., 2021.)

0 long (constitutes most of the length of the lower jaw)

1 short (constitutes less than half of jaw length)

**80. Teeth on dentary**

(Cloutier & Arratia 2004, Xu et al. 2014; Giles et al., 2017; López-Arbarello & Sferco 2018; Argyriou et al., 2018; Latimer & Giles, 2018; Wilson et al., 2019; Figueroa et al., 2021.)

0 present

1 absent

**81. [CH 22; G 67] Dentary with conspicuously reflexed distal tip**

(Friedman & Blom, 2006; Long et al., 2008; Swartz, 2009; Choo, 2011; Giles et al., 2015b; Giles et al., 2017; Argyriou et al., 2018; Latimer & Giles, 2018; Wilson et al., 2019; Figueroa et al., 2021.)

0 absent

1 present

**82. [CH 24; G 68] Enlarged series of parasymphysial teeth on dentary**

(Friedman & Blom, 2006; Long et al., 2008; Swartz, 2009; Choo, 2011; Giles et al., 2017; Argyriou et al., 2018; Latimer & Giles, 2018; Wilson et al., 2019; Figueroa et al., 2021.)

0 absent

1 present

**83. [CH 73; G69] Facet for parasymphysial tooth-whorl on anterior dentary**

(Choo, 2011; Giles et al., 2015b; Giles et al., 2017; Argyriou et al., 2018; Latimer & Giles, 2018; Wilson et al., 2019; Figueroa et al., 2021.)

0 present

1 absent

**84. [G 70] Teeth of outer dental arcade**

(Friedman, 2007; Giles et al., 2015b; Poyato-Ariza 2015; Giles et al., 2017; López-Arbarello & Sferco 2018; Argyriou et al., 2018; Latimer & Giles, 2018; Wilson et al., 2019; Figueroa et al., 2021. Coates (1998) states that the maxilla of *Melanecta* bears large teeth interspersed with smaller teeth, but it is unclear how these teeth are arranged. As such, this taxon is coded '?'. )

0 several rows of disorganized teeth

1 two rows, with large teeth lingually and small teeth labially

2 single row of teeth

**85. Jaw margins overlain by lateral lamina**

(Giles et al. 2017; Argyriou et al., 2018; Latimer & Giles, 2018; Wilson et al., 2019; Figueroa et al., 2021. In *Styracopterus*, *Fouldenia* and *Amphicentrum*, a lateral lamina of bone obscures the maxillary dentition (Sallan & Coates 2013).)

0 absent

1 present

**86. [CH 25; G 71] Acrodin caps on teeth**

(Patterson, 1982; Gardiner, 1984; Maisey, 1986; Gardiner & Schaeffer, 1989; Cloutier & Ahlberg, 1996; Taverne, 1997; Coates, 1999; Poplin & Lund, 2000; Schultze & Cumbaa, 2001; Zhu & Schultze, 2001; Zhu et al., 2001; Zhu & Yu, 2002; Cloutier & Arratia, 2004; Gardiner et al., 2005; Friedman & Blom, 2006; Zhu et al., 2006; Friedman, 2007; Long et al., 2008; Zhu et al., 2009; Friedman & Brazeau, 2010; Choo, 2011; Xu & Gao, 2011; Zhu et al., 2013; Xu et al., 2014; Giles et al., 2015b,c; Giles et al., 2017; Argyriou et al., 2018; Latimer & Giles, 2018; Wilson et al., 2019; Figueroa et al., 2021. )

0 absent

1 present

**87. Plicidentine**

(Zhu & Yu 2002; Friedman 2007; López-Arbarello 2012; Bermúdez-Rochas & Poyato-Ariza 2015; Poyato-Ariza 2015; Thies & Waschkewitz, 2015; Gibson 2016; Giles et al., 2017; López-Arbarello & Sferco 2018; Argyriou et al., 2018; Latimer & Giles, 2018; Wilson et al., 2019; Figueroa et al., 2021.)

0 absent

1 present

**88. [CH 27; G 73] Ossification of mentomeckelian region:**

(Friedman & Blom, 2006; Long et al., 2008; Swartz, 2009; Grande 2010; Choo, 2011; Xu et al., 2014; Giles et al., 2015b; Poyato-Ariza 2015; Giles et al., 2017; Argyriou et al., 2018; Latimer & Giles, 2018; Wilson et al., 2019; Figueroa et al., 2021.)

0 present

1 absent

**89. [CH 23; G 76] Number of infradentaries**

(Friedman & Blom, 2006; Friedman, 2007; Long et al., 2008; Choo, 2011; Xu & Gao, 2011; Xu et al., 2014; Giles et al., 2015b; Poyato-Ariza 2015; Giles et al., 2017; López-Arbarello & Sferco 2018; Argyriou et al., 2018; Latimer & Giles, 2018; Wilson et al., 2019; Figueroa et al., 2021.)

0 more than two

1 two (angular and surangular)

2 one (angular only)

**90. [G 74] Coronoids (sensu stricto, excluding parasymphysial tooth whorl or anterior coronoid)**

(Schultze and Cumbaa, 2001; Zhu and Schultze, 2001; Zhu et al., 2001; Zhu and Yu, 2002; Zhu et al., 2006; Friedman, 2007; Zhu et al., 2009; Giles et al., 2015b,c; Poyato-Ariza 2015; Giles et al., 2017; López-Arbarello & Sferco 2018; Argyriou et al., 2018; Latimer & Giles, 2018; Wilson et al., 2019; Figueroa et al., 2021. )

0 present

1 absent

**91. [G 75] Number of coronoids**

(Ahlberg & Clack, 1998; Daeschler et al., 2006; Long et al., 2006; Friedman, 2007; Zhu et al., 2009; Zhu et al., 2013; Giles et al., 2015b; Giles et al., 2017; Argyriou et al., 2018; Latimer & Giles, 2018; Wilson et al., 2019; Figueroa et al., 2021. A single specimen of *Pteronisculus* *stensioi* has at least five or six coronoids anterior to the prearticular region. However, these appear to correspond to the three coronoids present in most specimens, so the taxon is coded here as '2'. Two coronoids are reported in *Boreosomus* (Nielsen, 1942).)

0 five

1 four or more

2 three

3 two

4 one

**92. [G 76] Posterior coronoid**

(Cloutier & Ahlberg, 1996; Ahlberg & Johanson, 1998; Zhu & Ahlberg, 2004; Daeschler et al., 2006; Long et al., 2006; Giles et al., 2015b; Giles et al., 2017; Argyriou et al., 2018; Latimer & Giles, 2018; Wilson et al., 2019; Figueroa et al., 2021.)

0 morphologically similar to anterior coronoids

1 expanded

**93. [G 78] Coronoid process of lower jaw**

(Patterson 1973; Gardiner & Schaeffer 1989; Olsen & McCune 1991; Poyato-Ariza & Wenz, 2002; Zhu & Yu, 2002; Friedman, 2007; Friedman, 2007; Xu & Gao 2011; Xu et al. 2014, 2015; Giles et al., 2015b; Poyato-Ariza 2015; Xu & Zhao, 2016; Giles et al., 2017; Cawley & Kriwet 2018; Cawley & Kriwet 2018; López-Arbarello & Sferco 2018; Argyriou et al., 2018; Latimer & Giles, 2018; Wilson et al., 2019; Figueroa et al., 2021.)

0 absent

1 present

**94. Coronoid process contributed to by**

(Modified from Patterson 1973; Olsen & McCune 1991; Gardiner et al. 2005; Poyato-Ariza 2015; Giles et al., 2017; López-Arbarello & Sferco 2018; Argyriou et al., 2018; Latimer & Giles, 2018; Wilson et al., 2019; Figueroa et al., 2021.)

0 prearticular only

1 surangular only

2 dentary plus postdentary bones

3 angular only

**95. Leptolepid notch**

(Arratia 2013; Giles et al., 2017; López-Arbarello & Sferco 2018; Argyriou et al., 2018; Latimer & Giles, 2018; Wilson et al., 2019; Figueroa et al., 2021. A distinct notch in the posterior margin of the dentary is seen in taxa such as *Leptolepis*.)

0 absent

1 present

**96. Symplectic involvement in jaw joint**

(Grande & Bemis 1998; Olsen & McCune 1991; Grande 2010; Xu & Wu 2012; Xu et al. 2014, 2015; López-Arbarello 2012; Bermúdez-Rochas & Poyato-Ariza 2015; Poyato-Ariza 2015; Thies & Waschkewitz, 2015; Gibson 2016; Xu & Zhao, 2016; Giles et al., 2017; López-Arbarello & Sferco 2018; Argyriou et al., 2018; Latimer & Giles, 2018; Wilson et al., 2019; Figueroa et al., 2021.)

0 absent

1 present

**97. [G 79] Retroarticular process**

(Friedman, 2007; Giles et al., 2015b; Giles et al., 2017; Argyriou et al., 2018; Latimer & Giles, 2018; Wilson et al., 2019; Figueroa et al., 2021.)

0 present

1 absent

**98. Palatal bite**

(Giles et al., 2017; Argyriou et al., 2018; Latimer & Giles, 2018; Wilson et al., 2019; Figueroa et al., 2021.)

0 absent

1 present

**99. [G 82] Palatal articulation with basipterygoid process**

(Revised from Friedman, 2007; Brazeau, 2009; Zhu et al., 2009; Friedman & Brazeau, 2010; Davis et al., 2012; Zhu et al., 2013; Giles et al., 2015b,c; Giles et al., 2017; López-Arbarello & Sferco 2018; Argyriou et al., 2018; Latimer & Giles, 2018; Wilson et al., 2019; Figueroa et al., 2021. This character is expanded from previous formulations, which only considered whether a basipterygoid fenestra was absent or present. Where a basipterygoid process is absent, the dorsal margin of the palate may be flat, or the metapterygoid may bear a distinct notch.)

0 articulation not obvious

1 via basipterygoid fenestra

2 via metapterygoid process/notch

**100. Palatoquadrate ossifications**

(Giles et al., 2017; Argyriou et al., 2018; Latimer & Giles, 2018; Wilson et al., 2019; Figueroa et al., 2021.)

0 comineralized

1 separate ossification centers

**101. Lateral process of ectopterygoid**

(Giles et al., 2017; Argyriou et al., 2018; Latimer & Giles, 2018; Wilson et al., 2019; Figueroa et al., 2021.)

0 absent

1 present

**102. Palatoquadrate symphysis**

(Giles et al., 2017; Argyriou et al., 2018; Latimer & Giles, 2018; Wilson et al., 2019; Figueroa et al., 2021. This character captures whether the palatoquadrates contact at the midline. )

0 absent

1 present

**103. Dorsal margin of palate**

(Giles et al., 2017; Argyriou et al., 2018; Latimer & Giles, 2018; Wilson et al., 2019; Figueroa et al., 2021.)

0 high posterior extension

1 flat dorsal margin

**104. Metapterygoid posterior to quadrate**

(Giles et al., 2017; Argyriou et al., 2018; Latimer & Giles, 2018; Wilson et al., 2019; Figueroa et al., 2021.)

0 absent

1 present

**105. [G 81] Number of dermopalatines**

(Friedman, 2007; Giles et al., 2015b; Giles et al., 2017; Argyriou et al., 2018; Latimer & Giles, 2018; Wilson et al., 2019; Figueroa et al., 2021.)

0 multiple

1 single

**106. Prearticular**

(Poyato-Ariza 2015; Giles et al., 2017; López-Arbarello & Sferco 2018; Argyriou et al., 2018; Latimer & Giles, 2018; Wilson et al., 2019; Figueroa et al., 2021.)

0 present

1 absent

**107. Vomers**

(Patterson 1973; Olsen & McCune 1991; López-Arbarello 2012; Arratia 2013; Xu & Wu, 2012; Bermúdez-Rochas & Poyato-Ariza 2015; Poyato-Ariza 2015; Thies & Waschkewitz, 2015; Gibson 2016; Xu & Zhao, 2016; Giles et al., 2017; López-Arbarello & Sferco 2018; Argyriou et al., 2018; Latimer & Giles, 2018; Wilson et al., 2019; Figueroa et al., 2021.)

0 paired

1 single

**108. Vomer sutured to parasphenoid**

(Patterson 1973; Olsen & McCune 1991; Hurley et al. 2007; Giles et al., 2017; Argyriou et al., 2018; Latimer & Giles, 2018; Wilson et al., 2019; Figueroa et al., 2021.)

0 absent

1 present

**109. [CH 19; G 50] Accessory operculum**

(Schultze & Cumbaa, 2001; Cloutier & Arratia, 2004; Friedman & Blom, 2006; Long et al., 2008; Swartz, 2009; Giles et al., 2015b; Giles et al., 2017; Argyriou et al., 2018; Latimer & Giles, 2018; Wilson et al., 2019; Figueroa et al., 2021. This region of the cheek was removed in *Coccocephalichthys* (Poplin, 1974; Poplin & Véran, 1996), presumably by Watson (1925) when he first described the specimen. It is unclear from the surviving cast whether a dermohyal and/or accessory operculum were present, and as such this taxon is coded as ???.)

0 absent

1 present

**110. [CH 67; G 51] Operculum - relative size**

(Modified from Lund et al., 1995; Lund, 2000; Lund & Poplin, 2002; Cloutier & Arratia, 2004; Long et al., 2008; Swartz, 2009; Choo, 2011; López-Arbarello 2012; Bermúdez-Rochas & Poyato-Ariza 2015; Xu et al., 2015; Giles et al., 2015b; Thies & Waschkewitz, 2015; Gibson 2016; Xu & Zhao, 2016; Giles et al., 2017; López-Arbarello & Sferco 2018; Argyriou et al., 2018; Latimer & Giles, 2018; Wilson et al., 2019; Figueroa et al., 2021.)

0 at least twice as high as suboperculum

1 subequal

2 smaller than suboperculum

**111. Subopercle**

(Poyato-Ariza & Wenz, 2002; Xu et al. 2014; Giles et al., 2017; Cawley & Kriwet 2018; Argyriou et al., 2018; Latimer & Giles, 2018; Wilson et al., 2019; Figueroa et al., 2021.)

0 present

1 absent

**112. [CH 68; G 52] Anterodorsal process of suboperculum**

(Long et al., 2008; Choo, 2011; López-Arbarello 2012; Bermúdez-Rochas & Poyato-Ariza 2015; Giles et al., 2015b; Thies & Waschkewitz, 2015; Poyato-Ariza 2015; Gibson 2016; Giles et al., 2017; López-Arbarello & Sferco 2018; Argyriou et al., 2018; Latimer & Giles, 2018; Wilson et al., 2019; Figueroa et al., 2021. An anterodorsal process is present some non-neopterygian actinopterygians and 'semionotiforms'. Although these processes do not appear to be homologous they are coded within the character.)

0 absent

1 present

**113. Anteroventral process of suboperculum**

(Giles et al., 2017; Argyriou et al., 2018; Latimer & Giles, 2018; Wilson et al., 2019; Figueroa et al., 2021.)

0 absent

1 present

**114. [G 62] Number of cheek bones bearing pre-opercular canal posterior to jugal**

(Friedman, 2007; Zhu et al., 2009; Zhu et al., 2013; Giles et al., 2015b; Xu & Zhao, 2016; Giles et al., 2017; Argyriou et al., 2018; Latimer & Giles, 2018; Wilson et al., 2019; Figueroa et al., 2021.)

0 one

1 multiple

2 series of small ossicles

**115. Preoperculum orientation**

(Modified from Patterson 1973; Olsen & McCune 1991; Grande & Bemis 1998; Gardiner et al., 2005; Swartz 2009; López-Arbarello 2012; Bermúdez-Rochas & Poyato-Ariza 2015; Thies & Waschkewitz, 2015; Poyato-Ariza 2015; Gibson 2016; Giles et al., 2017; López-Arbarello & Sferco 2018; Argyriou et al., 2018; Latimer & Giles, 2018; Wilson et al., 2019; Figueroa et al., 2021. This character is reformulated from its original (compound) formulation, which considered both maxilla and preoperculum shape. Primitively in actinopterygians the preoperculum is wholly or partially developed dorsal to the maxilla as an anterodorsal-posteroventrally oriented bone, either with (e.g. *Mimipiscis, Moythomasia*) or without (e.g. *Cheirolepis*) a dorsoventrally oriented limb. The preoperculum may also be near-vertical, with no distinct anterodorsal or anteroventral extensions (e.g. *Boreosomus, Peltopleurus*), or developed as an anteroventral-posterodorsally-directed bone largely ventral to the maxilla (e.g. *Discoserra, Propterus*).)

0 prounounced dorsal limb

1 vertical

2 pronounced ventral limb

**116. Junction between preopercular and more anterior cheek bones**

(Modified from López-Arbarello 2012; Bermúdez-Rochas & Poyato-Ariza 2015; Poyato-Ariza 2015; Thies & Waschkewitz, 2015; Gibson 2016; Giles et al., 2017; López-Arbarello & Sferco 2018; Argyriou et al., 2018; Latimer & Giles, 2018; Wilson et al., 2019; Figueroa et al., 2021.)

0 Infraorbitals (including jugal) or suborbitals suture with or abut preopercular

1 Infraorbitals (including jugals) and suborbitals broadly overlap preopercular

2 completely obscure

**117. Posterior border of preopeculum notched ventrally**

(López-Arbarello 2012; Bermúdez-Rochas & Poyato-Ariza 2015; Giles et al., 2017; Thies & Waschkewitz, 2015; Gibson 2016; López-Arbarello & Sferco 2018; Argyriou et al., 2018; Latimer & Giles, 2018; Wilson et al., 2019; Figueroa et al., 2021.)

0 absent

1 present

**118. Interopercle**

(Patterson 1973; Grande & Bemis 1998; Gardiner & Schaeffer 1989; Olsen & McCune 1991; Poyato-Ariza & Wenz, 2002; Xu & Gao 2011; Xu et al. 2014, 2015. Gardiner & Schaeffer 1989; Olsen & McCune 1991; Gardiner et al. 1996; Gardiner et al. 2005; Cavin & Suteethorn 2006; Hurley et al. 2007; López-Arbarello 2012; Bermúdez-Rochas & Poyato-Ariza 2015; Poyato-Ariza 2015; Thies & Waschkewitz, 2015; Gibson 2016; Xu & Zhao, 2016; Giles et al., 2017; Cawley & Kriwet 2018; López-Arbarello & Sferco 2018; Argyriou et al., 2018; Latimer & Giles, 2018; Wilson et al., 2019; Figueroa et al., 2021. )

0 absent

1 present

**119. [CH 72; G 53] Branchiostegal rays - dorsal-most in series**

(Lund et al., 1995; Cloutier & Arratia, 2004; Choo, 2011; Giles et al., 2015b; Giles et al., 2017; Argyriou et al., 2018; Latimer & Giles, 2018; Wilson et al., 2019; Figueroa et al., 2021.)

0 of similar depth to adjacent branchiostegal ray

1 deeper than adjacent branchiostegal ray

**120. Lateral gulars**

(Olsen & McCune 1991; Xu et al., 2014; Giles et al., 2017; Argyriou et al., 2018; Latimer & Giles, 2018; Wilson et al., 2019; Figueroa et al., 2021. )

0 present

1 absent

**121. [CH 20; G 54] Lateral gulars**

(Gardiner & Schaeffer, 1989; Cloutier & Ahlberg, 1996; Taverne, 1997; Lund & Poplin, 1997; Coates, 1999; Schultze & Cumbaa, 2001; Zhu & Schultze, 2001; Cloutier & Arratia, 2004; Friedman & Blom, 2006; Long et al., 2008; Swartz, 2009; Brazeau, 2009; Xu & Gao, 2011; Davis et al., 2012; Zhu et al., 2013; Xu et al., 2014; Giles et al., 2015b,c; Giles et al., 2017; Argyriou et al., 2018; Latimer & Giles, 2018; Wilson et al., 2019; Figueroa et al., 2021. The condition in *Boreosomus* (Nielsen, 1942) is unique: instead of lateral gulars flanking a median gular, there appears to be a second median gular. This may well represent a fusion of the two, longer lateral gulars, is coded as such.)

0 extending most of the length of the lower jaw

1 restricted to the anterior third of the lower jaw (no longer than the width of three branchiostegals)

**122. [G 55] Median gular**

(Olsen & McCune 1991; Lund et al., 1995; Cloutier & Ahlberg, 1996; Coates, 1999; Lund, 2000; Schultze & Cumbaa, 2001; Zhu & Schultze, 2001; Zhu et al., 2001: Lund & Poplin, 2002; Zhu & Yu, 2002; Cloutier & Arratia, 2004; Zhu et al., 2006; Friedman, 2007; Zhu et al., 2009; Xu & Gao, 2011; López-Arbarello 2012; Zhu et al., 2013; Bermúdez-Rochas & Poyato-Ariza 2015; Poyato-Ariza 2015; Xu et al., 2014, 2015; Giles et al., 2015b,c; Thies & Waschkewitz, 2015; Xu & Zhao, 2015; Gibson 2016; Giles et al., 2017; López-Arbarello & Sferco 2018; Argyriou et al., 2018; Latimer & Giles, 2018; Wilson et al., 2019; Figueroa et al., 2021. Pearson & Westoll (1979: p. 365) state that a median gular is not known in *Cheirolepis canadensis*. Although a median gular is reconstructed by Cloutier & Arratia (1996: fig. 7), this bone is not present in any specimen photos and is not mentioned in the text. As such, this taxon is coded as ???.)

0 absent

1 present

**123. [G 56] Relative length of median gular**

(Giles et al., 2015b,c; Giles et al., 2017; Argyriou et al., 2018; Latimer & Giles, 2018; Wilson et al., 2019; Figueroa et al., 2021. The condition in *Boreosomus* (Nielsen, 1942) is unique: instead of lateral gulars flanking a median gular, there appears to be a second median gular. This may well represent a fusion of the two, longer lateral gulars, and is coded as such.)

0 much shorter than jaw length

1 more than half of law length

**124. [G 83] Fenestra ventrolateralis**

(Schultze & Cumbaa, 2001; Zhu & Schultze, 2001; Zhu et al., 2001; Zhu & Yu, 2002; Zhu et al., 2006; Friedman, 2007; Zhu et al., 2009; Zhu et al., 2013; Giles et al., 2015b; Giles et al., 2017; Argyriou et al., 2018; Latimer & Giles, 2018; Wilson et al., 2019; Figueroa et al., 2021. )

0 absent

1 present

**125. [G 84] Palatal opening surrounded by premaxilla, maxilla, dermopalatine and vomer (choana)**

(Zhu & Yu, 2002; Friedman, 2007; Giles et al., 2015b; Giles et al., 2017; Argyriou et al., 2018; Latimer & Giles, 2018; Wilson et al., 2019; Figueroa et al., 2021. Character can only be coded in taxa which possess all of these bones.)

0 absent

1 present

**126. [G 85] Internasal cavity**

(Ahlberg & Johanson, 1998; Zhu & Yu, 2002; Zhu & Ahlberg, 2004; Daeschler et al., 2006; Long et al., 2006; Friedman, 2007; Zhu et al., 2009; Zhu et al., 2013; Giles et al., 2015b,c; Giles et al., 2017; Argyriou et al., 2018; Latimer & Giles, 2018; Wilson et al., 2019; Figueroa et al., 2021.)

0 absent

1 present

**127. [G 86] Interorbital septum**

(Friedman, 2007; Zhu et al., 2009; Brazeau, 2009; Friedman & Brazeau, 2010; Davis et al., 2012; Zhu et al., 2013; Giles et al., 2015b,c; Giles et al., 2017; Argyriou et al., 2018; Latimer & Giles, 2018; Wilson et al., 2019; Figueroa et al., 2021. )

0 broad

1 narrow

**128. Optic foramen**

(Giles et al., 2017; Argyriou et al., 2018; Latimer & Giles, 2018; Wilson et al., 2019; Figueroa et al., 2021. Primitively in actinopts, the optic nerve exits the cranial cavity into the orbit through paired foramina approximately halfway up the orbital wall. In many Carboniferous and younger taxa, much of the orbital wall is unossified (the optic fenestra). The optic nerves may exit through openings just posterior to (e.g. *Pteronisculus*) or confluent with (e.g. *Pholidophorus*) the optic fenestra. In polypterids and *Fukangichthys*, the optic nerve exits ventrally through paired foramina that abut the parasphenoid.)

0 dorsally positioned

1 ventrally positioned (i.e. abuts parasphenoid)

**129. [G 87] Pronounced median anterior crista on dorsal surface of braincase**

(Giles et al., 2015b,c; Giles et al., 2017; Argyriou et al., 2018; Latimer & Giles, 2018; Wilson et al., 2019; Figueroa et al., 2021. Carboniferous and younger actinopts such as *Lawrenciella* (Hamel & Poplin, 2008) have a median crista anterior to the anterior dorsal fontanelle upon which the skull roof sits.)

0 absent

1 present

**130. [G 88] Expanded anterior dorsal fontanelle**

(Giles et al., 2015b,c; Giles et al., 2017; Argyriou et al., 2018; Latimer & Giles, 2018; Wilson et al., 2019; Figueroa et al., 2021. The anterior dorsal fontanelle of many Carboniferous and younger actinopts is greatly expanded, in contrast to the smaller fontanelle of Devonian taxa such as *Mimipiscis* (Gardiner, 1984).)

0 absent

1 present

**131. [G 89] Endoskeletal intracranial joint**

(Cloutier & Ahlberg, 1996; Ahlberg & Johanson, 1998; Zhu & Ahlberg, 2004; Zhu et al., 2001; Zhu & Yu, 2002; Daeschler et al., 2006; Long et al., 2006; Friedman, 2007; Brazeau, 2009; Zhu et al., 2009; Friedman & Brazeau, 2010; Davis et al., 2013; Zhu et al., 2013; Giles et al., 2015b,c; Giles et al., 2017; Argyriou et al., 2018; Latimer & Giles, 2018; Wilson et al., 2019; Figueroa et al., 2021.)

0 absent

1 present

**132. [G 90] Eye stalk or unfinished area for similar structure**

(Zhu & Schultze, 2001; Zhu et al., 2001; Zhu & Yu, 2002; Zhu et al., 2006; Friedman, 2007; Zhu et al., 2009; Zhu et al., 2013; Giles et al., 2015b,c; Giles et al., 2017; Argyriou et al., 2018; Latimer & Giles, 2018; Wilson et al., 2019; Figueroa et al., 2021. This character is coded as absent for taxa that possess a large interorbital fenestra (e.g. *Pteronisculus*, *Coccocephalichthys*, *Kentuckia deani*), as, if present, the eyestalk area would be visible posterior to the opening for the optic nerve.)

0 absent

1 present

**133. [G 91] Roof of posterior myodome perforated by palatine branch of facial nerve (VII)** (Coates, 1999; Giles et al., 2015b; Giles et al., 2017; Argyriou et al., 2018; Latimer & Giles, 2018; Wilson et al., 2019; Figueroa et al., 2021.)

0 absent

1 present

**134. [G 92] Foramen for abducens nerve (VI) dorsally positioned (level with optic foramen (II))**

(Coates, 1999; Giles et al., 2015b; Giles et al., 2017; Argyriou et al., 2018; Latimer & Giles, 2018; Wilson et al., 2019; Figueroa et al., 2021.)

0 absent

1 present

**135. [G 93] Anterodorsal myodome**

(Patterson 1973; Olsen & McCune 1991; Coates, 1999; Gardiner et al. 1996; Hurley et al. 2007; Xu & Gao, 2011; Xu et al., 2014, 2015; Giles et al., 2015b; Poyato-Ariza 2015; Xu & Zhao, 2016; Giles et al., 2017; Argyriou et al., 2018; Latimer & Giles, 2018; Wilson et al., 2019; Figueroa et al., 2021.)

0 paired

1 single

2 absent

**136. [G 94] Posterior myodome**

(Modified from Patterson 1973; Olsen & McCune 1991; Coates, 1999; Xu et al., 2014. Wiley 1976; Gardiner, 1984; Gardiner & Schaeffer, 1989; Gardiner et al. 1996; Hurley et al. 2007; López-Arbarello 2012; Xu & Gao, 2011; Bermúdez-Rochas & Poyato-Ariza 2015; Xu et al. 2014, 2015; Giles et al., 2015b; Thies & Waschkewitz, 2015; Gibson 2016; Xu & Zhao, 2016; Giles et al., 2017; López-Arbarello & Sferco 2018; Argyriou et al., 2018; Latimer & Giles, 2018; Wilson et al., 2019; Figueroa et al., 2021.)

0 absent

1 paired

2 median

**137. [G 96] Basicranial fenestra**

(Ahlberg & Johanson, 1998; Zhu et al., 2001; Zhu & Yu, 2002; Zhu & Ahlberg, 2004; Friedman, 2007; Zhu et al., 2009; Zhu et al., 2013; Giles et al., 2015b,c; Giles et al., 2017; Argyriou et al., 2018; Latimer & Giles, 2018; Wilson et al., 2019; Figueroa et al., 2021. )

0 absent

1 present

**138. [G 98] Endoskeletal spiracular canal**

(Modified from Patterson, 1982; Gardiner, 1984; Gardiner & Schaeffer, 1989; Taverne, 1997; Coates, 1999; Gardiner et al., 2005; Xu & Gao, 2011; Xu et al., 2014; Giles et al., 2015b; Giles et al., 2017; López-Arbarello & Sferco 2018; Argyriou et al., 2018; Latimer & Giles, 2018; Wilson et al., 2019; Figueroa et al., 2021. Taxa that lack a groove on the lateral commissure are coded as inapplicable for this character.) )

0 open

1 partial closure of spiracular bar

2 complete enclosure in canal

**139. [G 100] Basipterygoid process**

(Gardiner et al., 2005; Xu & Gao, 2011; Xu et al., 2014, 2015; Giles et al., 2015b; Xu & Zhao, 2016; Giles et al., 2017; López-Arbarello & Sferco 2018; Argyriou et al., 2018; Latimer & Giles, 2018; Wilson et al., 2019; Figueroa et al., 2021.)

0 present

1 absent

**140. [G 101] Basipterygoid process with vertically oriented component**

(Ahlberg & Johanson, 1998; Zhu & Schultze, 2001; Zhu et al., 2001; Zhu & Yu, 2002; Zhu & Ahlberg, 2004; Zhu et al., 2006; Friedman, 2007; Zhu et al., 2009; Davis et al., 2012; Zhu et al., 2013; Giles et al., 2015b,c; Giles et al., 2017; Argyriou et al., 2018; Latimer & Giles, 2018; Wilson et al., 2019; Figueroa et al., 2021.)

0 absent

1 present

**141. [G 103] Dermal component to basipterygoid process**

(Gardiner, 1984; Gardiner & Schaeffer, 1989; Taverne, 1997; Coates, 1999; Giles et al., 2015b; Giles et al., 2017; Argyriou et al., 2018; Latimer & Giles, 2018; Wilson et al., 2019; Figueroa et al., 2021.)

0 absent

1 present

**142. Hyoid facet**

(Gardiner et al. 2005; Gardiner et al. 1996; Hurley et al. 2007; Xu & Gao 2011; Xu et al. 2015; Giles et al., 2017; López-Arbarello & Sferco 2018; Argyriou et al., 2018; Latimer & Giles, 2018; Wilson et al., 2019; Figueroa et al., 2021.)

0 directed posteroventrally

1 horizontal

**143. [G 104] Fossa bridgei**

(Gardiner, 1984; Gardiner & Schaeffer, 1989; Taverne, 1997; Coates, 1999; Xu & Gao, 2011; Xu et al., 2014; Giles et al., 2015b; Giles et al., 2017; Argyriou et al., 2018; Latimer & Giles, 2018; Wilson et al., 2019; Figueroa et al., 2021.)

0 absent

1 present

**144. [G 105] Posttemporal fossae**

(Zhu & Yu, 2002; Friedman, 2007; Argyriou et al., 2018; Latimer & Giles, 2018; Wilson et al., 2019; Figueroa et al., 2021.)

0 absent

1 present

**145. [G 106] Vestibular fontanelle**

(Friedman, 2007; Brazeau, 2009; Zhu et al., 2009; Friedman & Brazeau, 2010; Davis et al., 2012; Zhu et al., 2013; Brazeau & Friedman, 2014; Giles et al., 2015b,c; Giles et al., 2017; Argyriou et al., 2018; Latimer & Giles, 2018; Wilson et al., 2019; Figueroa et al., 2021.)

0 absent

1 present

**146. [G 107] Ventral cranial fissure and vestibular fontanelle**

(Coates, 1999; Giles et al., 2015b; Giles et al., 2017; Argyriou et al., 2018; Latimer & Giles, 2018; Wilson et al., 2019; Figueroa et al., 2021. We follow Coates (1999) in coding *Howqualepis* as '0' on the basis of Long 1988 fig. 16 and AMF65495 (pers. obs. S.G.), rather than the braincase reconstruction (Long, 1988: fig. 18.)

0 separated by bridge of bone

1 conflUent

**147. [G 108] Accessory fenestration in otic capsule**

(Friedman, 2007; Zhu et al., 2009; Giles et al., 2015b; Giles et al., 2017; Argyriou et al., 2018; Latimer & Giles, 2018; Wilson et al., 2019; Figueroa et al., 2021.)

0 absent

1 present

**148. [G 109] Otoccipital fissure**

(Friedman, 2007; Brazeau, 2009; Davis et al., 2012; Zhu et al., 2013; Giles et al., 2015b,c; Giles et al., 2017; Argyriou et al., 2018; Latimer & Giles, 2018; Wilson et al., 2019; Figueroa et al., 2021.)

0 absent

1 present

**149. [G 110] Median projection overhanging posterior part of posterior dorsal fontanelle**

(Giles et al., 2015b; Giles et al., 2017; Argyriou et al., 2018; Latimer & Giles, 2018; Wilson et al., 2019; Figueroa et al., 2021. Variable in *Boreosomus*: the posterior dorsal fontanelle is closed in the specimen figured in Nielsen (1942: plate 25F), but developed in the specimen figured in plate 28. This taxon is coded ?0/1? to reflect this polymorphism.)

0 absent

1 present

**150. [G 111] Median projection overhanging anterior part of posterior dorsal fontanelle**

(Giles et al., 2015b; Giles et al., 2017; Argyriou et al., 2018; Latimer & Giles, 2018; Wilson et al., 2019; Figueroa et al., 2021. This projection is somewhat reduced in *Pteronisculus* (Nielsen, 1942), but is coded '1' here. Variable in *Boreosomus*: the posterior dorsal fontanelle is closed in the specimen figured in Nielsen 1942 plate 25F, but developed in the specimen figured in plate 28. This taxon is coded ?0/1? to reflect this polymorphism. *Cheirolepis trailli* is coded '0' (2015a).)

0 absent

1 present

**151. [G 112] Dorsal aorta**

(Coates & Sequeira, 1998; Coates & Sequeira, 2001a, b; Coates, 1999; Friedman, 2007; Zhu et al., 2009; Friedman & Brazeau, 2010; Zhu et al., 2013; Giles et al., 2015b,c; Giles et al., 2017; López-Arbarello & Sferco 2018; Argyriou et al., 2018; Latimer & Giles, 2018; Wilson et al., 2019; Figueroa et al., 2021. This character is coded as inapplicable in taxa that lack a canal for the dorsal aorta. *Cheirolepis trailli* is coded '0' (Giles et al., 2015a). The aortic canal of *Moythomasia* is not figured by Gardiner (1984), but a clear posterior notch in the aortic canal can be seen in Long & Trinajstic (2010:fig 5b). The neurocranium of *Gogosardina* is not yet described, but this character can be coded on the basis of Choo et al. (2009: fig. 9).)

0 open in groove

1 canal notched posteriorly

2 completely enclosed in canal

**152. [G 113] Dorsal aorta pierced by canal/s for exit of eff.a.2**

(Giles et al., 2015b; Giles et al., 2017; Argyriou et al., 2018; Latimer & Giles, 2018; Wilson et al., 2019; Figueroa et al., 2021. In *Mimipiscis bartrami* and *M. toombsi*, a groove for one of the efferent branchial arteries branches off from the lateral dorsal aorta immediately before the articular area for the first infrapharyngobranchial. However, it is uncertain which, so both taxa coded as '?' for these characters. The neurocranium of *Gogosardina* is not yet described, but this character can be coded on the basis of Choo et al. (2009: fig. 9).)

0 absent

1 present

**153. [G 114] Dorsal aorta pierced by canal/s for exit of eff.a.1**

(Giles et al., 2015b; Giles et al., 2017; Argyriou et al., 2018; Latimer & Giles, 2018; Wilson et al., 2019; Figueroa et al., 2021. In *Mimipiscis bartrami* and *M. toombsi*, a groove for one of the efferent branchial arteries branches off from the lateral dorsal aorta immediately before the articular area for the first infrapharyngobranchial. However, it is uncertain which, so both taxa coded as '?' for these characters. The neurocranium of *Gogosardina* is not yet described, but this character can be coded on the basis of Choo et al. (2009: fig. 9).)

0 absent

1 present

**154. [G 115] Bifurcation of dorsal aorta**

(Coates & Sequeira, 1998; Coates & Sequeira, 2001a, b; Coates, 1999; Friedman, 2007; Zhu et al., 2009; Friedman & Brazeau, 2010; Zhu et al., 2013; Giles et al., 2015b,c; Giles et al., 2017; Argyriou et al., 2018; Latimer & Giles, 2018; Wilson et al., 2019; Figueroa et al., 2021.)

0 posterior to occiput

1 anterior to occiput

**155. [G 116] Birfurcation of dorsal aorta into lateral dorsal aortae**

(Coates, 1999; Giles et al., 2015b; Giles et al., 2017; Argyriou et al., 2018; Latimer & Giles, 2018; Wilson et al., 2019; Figueroa et al., 2021. This character is coded as inapplicable in taxa that lack a canal for the dorsal aorta. In *Mimipiscis toombsi*, the bifurcation point of the dorsal aorta can be open (Giles & Friedman, 2014: fig. 2) or closed (Gardiner 1984: fig. 15). This taxon is coded '0/1' to reflect this polymorphism. The aortic canal of *Moythomasia* is not figured by Gardiner (1984), but the bifucation into the lateral dorsal aortae can be seen in Long & Trinajstic (2010:fig 5b).)

0 open

1 enclosed in canal

**156. Braincase ossifications differentiated**

(Giles et al., 2017; Argyriou et al., 2018; Latimer & Giles, 2018; Wilson et al., 2019; Figueroa et al., 2021.)

0 absent

1 present

**157. Basisphenoid**

(Wiley 1976; López-Arbarello 2012; Bermúdez-Rochas & Poyato-Ariza 2015; Thies & Waschkewitz, 2015; Gibson 2016; Giles et al., 2017; López-Arbarello & Sferco 2018; Argyriou et al., 2018; Latimer & Giles, 2018; Wilson et al., 2019; Figueroa et al., 2021.)

0 present

1 absent or very reduced

**158. Opisthotic-pterotic relationship**

(Gardiner et al. 1996; Hurley et al. 2007; Giles et al., 2017; Argyriou et al., 2018; Latimer & Giles, 2018; Wilson et al., 2019; Figueroa et al., 2021. This character can only be coded when separate braincase ossifications can be identified.)

0 opisthotic larger than subotic

1 opisthotic and pterotic equal in size

**159. Epioccipital**

(Hurley et al. 2007; Giles et al., 2017; Argyriou et al., 2018; Latimer & Giles, 2018; Wilson et al., 2019; Figueroa et al., 2021. This character can only be coded when separate braincase ossifications can be identified.)

0 present

1 absent

**160. Forward extension of the exoccipital around the vagus nerve**

(Olsen & McCune 1991; Gardiner et al. 1996; Cavin & Suteethorn 2006; Hurley et al. 2007; López-Arbarello 2012; Bermúdez-Rochas & Poyato-Ariza 2015; Thies & Waschkewitz, 2015; Gibson 2016; Giles et al. 2017; López-Arbarello & Sferco 2018; Argyriou et al., 2018; Latimer & Giles, 2018; Wilson et al., 2019; Figueroa et al., 2021. This character can only be coded when separate braincase ossifications can be identified.)

0 absent

1 present

**161. Spenotic with small dermal component**

(Grande 2010; López-Arbarello 2012; Xu & Wu 2012; Xu et al. 2014, 2015; Arratia 2013; Bermúdez-Rochas & Poyato-Ariza 2015; Poyato-Ariza 2015; Thies & Waschkewitz, 2015; Gibson 2016; Xu & Zhao, 2016; Giles et al. 2017; López-Arbarello & Sferco 2018; Argyriou et al., 2018; Latimer & Giles, 2018; Wilson et al., 2019; Figueroa et al., 2021. This character can only be coded when separate braincase ossifications can be identified.)

0 absent

1 present

**162. Pterotic**

(Gardiner et al. 1996; Grande & Bemis, 1998; Hurley et al. 2007; Xu et al. 2014; Poyato-Ariza 2015; Xu & Shen, 2015; Xu & Zhao, 2016; Giles et al. 2017; López-Arbarello & Sferco 2018; Argyriou et al., 2018; Latimer & Giles, 2018; Wilson et al., 2019; Figueroa et al., 2021. This character can only be coded when separate braincase ossifications can be identified.)

0 present

1 absent

**163. Opisthotic bone**

(Wiley 1976; Olsen & McCune 1991; Grande & Bemis 1998; Cavin & Suteethorn 2006; Hurley et al. 2007; Grande 2010; López-Arbarello 2012; Bermúdez-Rochas & Poyato-Ariza 2015; Poyato-Ariza 2015; Thies & Waschkewitz, 2015; Xu et al., 2014, 2015; Gibson 2016; Xu & Zhao, 2016; Giles et al. 2017; López-Arbarello & Sferco 2018; Argyriou et al., 2018; Latimer & Giles, 2018; Wilson et al., 2019; Figueroa et al., 2021. This character can only be coded when separate braincase ossifications can be identified.)

0 present

1 absent

**164. Intercalar**

(Olsen & McCune 1991; Olsen 1994; Gardiner et al. 1996; López-Arbarello 2012; Xu et al. 2014; Poyato-Ariza 2015; Xu & Shen, 2015; Thies & Waschkewitz, 2015; Gibson 2016; Xu & Zhao, 2016; Giles et al. 2017; López-Arbarello & Sferco 2018; Argyriou et al., 2018; Latimer & Giles, 2018; Wilson et al., 2019; Figueroa et al., 2021. This character can only be coded when separate braincase ossifications can be identified. )

0 present

1 absent

**165. Supraoccipital bone**

(Grande 2010; Poyato-Ariza 2015; Xu et al. 2014, 2015; Xu & Zhao, 2016; Giles et al. 2017; López-Arbarello & Sferco 2018; Argyriou et al., 2018; Latimer & Giles, 2018; Wilson et al., 2019; Figueroa et al., 2021. This character can only be coded when separate braincase ossifications can be identified. )

0 absent

1 present

**166. Membranous outgrowth of intercalar**

(Patterson 1973; Olsen & McCune 1991; Gardiner et al. 1996; Hurley et al. 2007; Giles et al. 2017; López-Arbarello & Sferco 2018; Argyriou et al., 2018; Latimer & Giles, 2018; Wilson et al., 2019; Figueroa et al., 2021. This character can only be coded when separate braincase ossifications can be identified. )

0 absent

1 present

**167. Post-temporal fossa**

(Patterson 1973; Gardiner, 1984; Gardiner & Schaeffer, 1989; Olsen & McCune 1991; Coates, 1999; Hurley et al. 2007; López-Arbarello 2012; Xu & Gao, 2011; Xu et al. 2014, 2015; Bermúdez-Rochas & Poyato-Ariza 2015; Giles et al., 2015b; Poyato-Ariza 2015; Thies & Waschkewitz, 2015; Gibson 2016; Xu & Zhao, 2016; Giles et al. 2017; Argyriou et al., 2018; Latimer & Giles, 2018; Wilson et al., 2019; Figueroa et al., 2021.)

0 absent

1 present

**168. Sub-temporal fossa**

(Gardiner, 1984; Gardiner & Schaeffer, 1989; Gardiner et al. 1996; Hurley et al. 2007; Xu & Gao, 2011; Xu et al. 2014, 2015; Xu & Zhao, 2016; Argyriou et al., 2018; Latimer & Giles, 2018; Wilson et al., 2019; Figueroa et al., 2021.)

0 absent

1 present

**169. Dilatator fossa**

(Gardiner, 1984; Gardiner & Schaeffer, 1989; Coates, 1999; Gardiner et al. 1996; Hurley et al. 2007; Xu & Gao, 2011; Xu et al. 2014, 2015; Xu & Zhao, 2016; Giles et al. 2017; Argyriou et al., 2018; Latimer & Giles, 2018; Wilson et al., 2019; Figueroa et al., 2021.)

0 absent

1 present

**170. [G 117] Parasphenoid**

(Gardiner, 1984; Brazeau, 2009; Davis et al., 2012; Zhu et al., 2013; Giles et al., 2015b,c; Giles et al. 2017; Argyriou et al., 2018; Latimer & Giles, 2018; Wilson et al., 2019; Figueroa et al., 2021.)

0 absent

1 present

**171. [G 118] Parasphenoid**

(Coates, 1999; Zhu & Yu, 2002; Gardiner et al., 2005; Friedman, 2007, Xu & Gao, 2011; Xu et al., 2014, 2015; Giles et al., 2015b; Xu & Zhao, 2016; Giles et al. 2017; López-Arbarello & Sferco 2018; Argyriou et al., 2018; Latimer & Giles, 2018; Wilson et al., 2019; Figueroa et al., 2021.)

0 terminates at/anterior to ventral otic fissure

1 extends across ventral otic fissure

2 extends to basioccipital

**172. [CH 28; G 120] Ascending process of the parasphenoid**

(Patterson, 1982; Coates, 1999; Dietze, 2000; Schultze & Cumbaa, 2001; Zhu & Schultze, 2001; Cloutier & Arratia, 2004; Gardiner et al., 2005; Friedman & Blom, 2006; Zhu et al., 2006; Zhu et al., 2009; Choo, 2011; Xu & Gao, 2011; Zhu et al., 2013; Xu et al., 2014; Giles et al., 2015b,c; Giles et al. 2017; López-Arbarello & Sferco 2018; Argyriou et al., 2018; Latimer & Giles, 2018; Wilson et al., 2019; Figueroa et al., 2021.)

0 absent

1 present

**173. [CH 29; G 121] Parasphenoid with multifid anterior margin**

(Friedman & Blom, 2006; Friedman, 2007; Zhu et al., 2009; Choo, 2011; Zhu et al., 2013; Giles et al., 2015b,c; Giles et al. 2017; Argyriou et al., 2018; Latimer & Giles, 2018; Wilson et al., 2019; Figueroa et al., 2021.)

0 absent

1 present

**174. [G 124] Buccohypophyseal canal pierces parasphenoid**

(Giles et al., 2015b,c; Giles et al. 2017; Argyriou et al., 2018; Latimer & Giles, 2018; Wilson et al., 2019; Figueroa et al., 2021. The buccohypophyseal canal typically enters the dorsal surface of the parasphenoid, but whether it exits via the ventral surface is more variable, and this distribution is captured by this character.)

0 present

1 absent

**175. Parasphenoid teeth**

(Arratia 2013; Giles et al. 2017; López-Arbarello & Sferco 2018; Argyriou et al., 2018; Latimer & Giles, 2018; Wilson et al., 2019; Figueroa et al., 2021.)

0 small

1 large

2 absent

**176. Parasphenoid pierced by internal carotid artery**

(Gardiner et al. 1996; Hurley et al. 2007; Xu & Wu, 2012; Xu et al., 2015; Xu & Zhao, 2016; Giles et al. 2017; Argyriou et al., 2018; Latimer & Giles, 2018; Wilson et al., 2019; Figueroa et al., 2021.)

0 absent

1 present

**177. Parasphenoid pierced by efferent pseudobranchial artery**

(Gardiner et al. 1996; Hurley et al. 2007; Xu & Wu, 2012; Xu et al., 2015; Xu & Zhao, 2016; Giles et al. 2017; Argyriou et al., 2018; Latimer & Giles, 2018; Wilson et al., 2019; Figueroa et al., 2021.)

0 absent

1 present

**178. Aortic notch in parasphenoid**

(Modified from Gardiner et al. 2005; Giles et al. 2017; Argyriou et al., 2018; Latimer & Giles, 2018; Wilson et al., 2019; Figueroa et al., 2021.)

0 absent

1 present

**179. Parabasal canal**

(Xu and Gao 2011; Xu et al. 2014; Giles et al. 2017; Argyriou et al., 2018; Latimer & Giles, 2018; Wilson et al., 2019; Figueroa et al., 2021.)

0 present

1 absent

**180. [G 125] Anterolaterally divergent olfactory tracts**

(Coates, 1999; Giles & Friedman, 2014; Giles et al., 2015b; Giles et al. 2017; Argyriou et al., 2018; Latimer & Giles, 2018; Wilson et al., 2019; Figueroa et al., 2021.)

0 absent

1 present

**181. [G 126] Elongate olfactory tract(s)**

(Brazeau, 2009; Friedman & Brazeau 2010; Davis et al., 2012; Zhu et al., 2013; Brazeau & Friedman, 2014; Giles & Friedman, 2014; Giles et al., 2015b,c; Giles et al. 2017; Argyriou et al., 2018; Latimer & Giles, 2018; Wilson et al., 2019; Figueroa et al., 2021. The olfactory tracts of *Osorioichthys* are elongate (pers. obs. unpubl. scan data S.G.).)

0 absent

1 present

**182. [G 127] Olfactory nerves carried in a single tract**

(Coates, 1999; Giles & Friedman, 2014; Giles et al., 2015b; Giles et al. 2017; Argyriou et al., 2018; Latimer & Giles, 2018; Wilson et al., 2019; Figueroa et al., 2021. The olfactory nerves are carried in separate tracts in *Osorioichthys* (pers. obs. unpubl. scan data S.G.).)

0 present

1 absent

**183. [G 128] Hypophyseal chamber**

(Coates, 1999; Xu & Gao, 2011; Xu et al., 2014; Giles et al., 2015b; Giles et al. 2017; Argyriou et al., 2018; Latimer & Giles, 2018; Wilson et al., 2019; Figueroa et al., 2021.)

0 projects posteroventrally

1 projects ventrally or anteroventrally

**184. [G 129] Optic lobes**

(Giles & Friedman, 2014; Giles et al., 2015b; Giles et al. 2017; Argyriou et al., 2018; Latimer & Giles, 2018; Wilson et al., 2019; Figueroa et al., 2021.)

0 narrower than cerebellum

1 same width or wider than cerebellum

**185. Optic lobes**

(Coates 1999; Hurley et al. 2007; Xu et al. 2014; Giles et al. 2017; Argyriou et al., 2018; Latimer & Giles, 2018; Wilson et al., 2019; Figueroa et al., 2021. )

0 smaller than telencephalon

1 larger than telencephalon

**186. [G 130] Optic tectum divided into bilateral halves**

(Coates, 1999; Giles et al., 2015b; Giles et al. 2017; Argyriou et al., 2018; Latimer & Giles, 2018; Wilson et al., 2019; Figueroa et al., 2021.)

0 absent

1 present

**187. [G 131] Cerebellar corpus**

(Giles et al., 2015b,c; Giles et al. 2017; Argyriou et al., 2018; Latimer & Giles, 2018; Wilson et al., 2019; Figueroa et al., 2021. The region posterior to the cerebellar auricles in *Lawrenciella* was considered to be the area octavolateralis by Hamel & Poplin (2005). However, we interpret it as the corpus cerebellum, and this taxon is coded '1'.)

0 absent

1 present

**188. Cerebellar corpus**

(Coates 1999; Hurley et al. 2007; Xu & Gao 2011; Xu et al. 2014; Giles et al. 2017; Argyriou et al., 2018; Latimer & Giles, 2018; Wilson et al., 2019; Figueroa et al., 2021. )

0 divided bilaterally

1 undivided

**189. Position of cerebellar corpus**

(Coates 1999; Hurley et al. 2007; Xu & Gao 2011; Xu et al. 2014; Giles et al. 2017; Argyriou et al., 2018; Latimer & Giles, 2018; Wilson et al., 2019; Figueroa et al., 2021. )

0 enters fourth ventricle

1 arches above fourth ventricle

**190. Cerebellar corpus with median anteriorly projecting portion**

(Coates 1999; Hurley et al. 2007; Xu & Gao 2011; Xu et al. 2014; Giles et al. 2017; Argyriou et al., 2018; Latimer & Giles, 2018; Wilson et al., 2019; Figueroa et al., 2021. )

0 absent

1 present

**191. [G 132] Horizontal semicircular canal**

(Davis et al., 2012; Zhu et al., 2013; Giles & Friedman, 2014; Giles et al., 2015b,c; Giles et al. 2017; Argyriou et al., 2018; Latimer & Giles, 2018; Wilson et al., 2019; Figueroa et al., 2021.)

0 joins vestibular region dorsal to ampulla for the posterior semicircular canal

1 joins vestibular region level with ampulla for the posterior semicircular canal

**192. [G 133] Junction between ampulla of posterior semicircular canal and cranial cavity**

(Giles et al., 2015b,c; Giles et al. 2017; Argyriou et al., 2018; Latimer & Giles, 2018; Wilson et al., 2019; Figueroa et al., 2021. In certain stem actinopterygians, such as *Mimipiscis* (Giles and Friedman, 2014), a short length of canal lies between the posterior ampulla and the remainder of the labyrinth.)

0 separated by short length of canal

1 confluent

**193. [G 134] Crus commune of anterior and posterior semicircular canal**

(Giles & Friedman, 2014; Giles et al., 2015b; Giles et al. 2017; Argyriou et al., 2018; Latimer & Giles, 2018; Wilson et al., 2019; Figueroa et al., 2021.)

0 dorsal to endocranial roof

1 ventral to endocranial roof

**194. [G 135] Lateral cranial canal**

(Gardiner, 1984; Gardiner & Schaeffer, 1989; Coates, 1999; Cloutier & Arratia, 2004; Gardiner et al., 2005; Zhu et al., 2006; Zhu et al., 2009; Zhu et al., 2013; Giles & Friedman, 2014; Xu et al., 2014; Giles et al., 2015b,c; Giles et al. 2017; Argyriou et al., 2018; Latimer & Giles, 2018; Wilson et al., 2019; Figueroa et al., 2021. The presence of a lateral cranial canal in *Ligulalepis* and *Psarolepis* is uncertain, but its presence in *Meemannia* is confirmed following Lu et al. (2016). *Erpetoichthys* is conservatively coded as '?'. )

0 absent

1 present

**195. Lateral cranial canal connects to cranial cavity anteriorly**

(Giles et al. 2017; Argyriou et al., 2018; Latimer & Giles, 2018; Wilson et al., 2019; Figueroa et al., 2021.)

0 absent

1 present

**196. [CH 30 in part; G 147] Enameloid on dermal bones and scales**

(Giles et al., 2015b,c, 2017; Argyriou et al., 2018; Latimer & Giles, 2018; Wilson et al., 2019; Figueroa et al., 2021: Characters 147-150 form part of an atomisation of the compound characters 'ganoine' (typically defined as a single or multilayer enamel covering) and 'cosmine' (typically defined as a single layer of enamel with a well-defined pore canal network) (e.g. Cloutier & Ahlberg, 1996; Ahlberg & Johanson, 1998; Zhu & Ahlberg, 2004; Schultze & Cumbaa, 2001; Zhu & Schultze, 2001; Zhu et al., 2001; Zhu & Yu, 2002; Daeschler et al., 2006; Long et al., 2006; Zhu et al., 2006; Zhu et al., 2009; Davis et al., 2012; Zhu et al., 2013). A similar approach to atomization was taken by Friedman (2007), Brazeau & Friedman (2010) and Giles et al. (2015b). As detailed histological investigations have not been carried out for the majority of early actinopterygians (rather, they have simply been described as being covered in/bearing ridges of ganoine), many of these characters cannot be coded for a number of taxa.)

0 absent

1 present

**197. [G 148] Extensive pore-canal network**

(See notes above for c. 196.)

0 absent

1 present

**198. [CH 30 in part; G 149] Enamel**

(See notes above for c. 196.)

0 single-layered

1 multi-layered

**199. [CH 30 in part; G 150] Enamel layers**

(See notes above for c. 196.)

0 applied directly to one another

1 separated by layers of dentine

**200. Scales on body**

(Giles et al. 2017; Argyriou et al., 2018; Latimer & Giles, 2018; Wilson et al., 2019; Figueroa et al., 2021.)

0 present

1 absent

**201. [G 151] Scales**

(Cloutier & Arratia, 2004; Friedman & Blom, 2006; Long et al., 2008; Swartz, 2009; Zhu et al., 2009; Choo, 2011; Giles et al., 2015b; Giles et al. 2017; Argyriou et al., 2018; Latimer & Giles, 2018; Wilson et al., 2019; Figueroa et al., 2021.)

0 micromeric

1 macromeric

**202. [CH 32 in part; G 152] Scales with 'peg and socket articulation'**

(Maisey, 1986; Gardiner & Schaeffer, 1989; Cloutier & Ahlberg, 1996; Coates, 1999; Dietze, 2000; Poplin & Lund, 2000; Schultze & Cumbaa, 2001; Cloutier & Arratia, 2004; Friedman & Blom, 2006; Friedman, 2007; Long et al., 2008; Brazeau, 2009; Swartz, 2009; Zhu et al., 2009; Friedman & Brazeau, 2010; López-Arbarello 2012; Xu & Gao, 2011; Choo, 2011; Davis et al., 2012; Zhu et al., 2013; Xu et al., 2014; Giles et al., 2015b,c; Xu & Zhao, 2016; Giles et al. 2017; López-Arbarello & Sferco 2018; Argyriou et al., 2018; Latimer & Giles, 2018; Wilson et al., 2019; Figueroa et al., 2021. This character is coded only for taxa that possess rhombic scales.)

0 absent

1 present

**203. [CH 32 in part; G 153] Peg on rhomboid scale**

(Patterson, 1982; Cloutier & Ahlberg, 1996; Dietze, 2000; Schultze & Cumbaa, 2001; Zhu & Schultze, 2001; Zhu et al., 2001: Zhu & Yu, 2002; Cloutier & Arratia, 2004; Friedman & Blom, 2006; Zhu et al., 2006; Friedman, 2007; Zhu et al., 2009; Giles et al. 2017; Argyriou et al., 2018; Latimer & Giles, 2018; Wilson et al., 2019; Figueroa et al., 2021. Although peg-and-socket articulation of is present between the scales of *Limnomis* (Daeschler, 2000), the nature of the peg is not described. As such, this taxon is conservatively coded '?'.)

0 narrow

1 broad

**204. [CH 33; G 154] Anterodorsal process on scale**

(Patterson, 1982; Gardiner, 1984; Gardiner & Schaeffer, 1989; Schultze & Cumbaa, 2001; Zhu & Schultze, 2001; Zhu et al., 2001; Zhu & Yu, 2002; Cloutier & Arratia, 2004; Friedman & Blom, 2006; Zhu et al., 2006; Friedman, 2007; Long et al., 2008; Swartz, 2009; Zhu et al., 2009; Choo, 2011; Zhu et al., 2013; Giles et al., 2015b,c; Giles et al. 2017; Argyriou et al., 2018; Latimer & Giles, 2018; Wilson et al., 2019; Figueroa et al., 2021.)

0 absent

1 present

**205. [CH 35; G 155] Scales with well developed pores on surface**

(Friedman & Blom 2006; Long et al., 2008; Swartz, 2009; Choo, 2011; Xu et al., 2014; Giles et al., 2015b; Giles et al. 2017; Argyriou et al., 2018; Latimer & Giles, 2018; Wilson et al., 2019; Figueroa et al., 2021. Scales from the posterior half of the flank in *Wendyichthys* bear pores on the enamel surface, whereas those from the anterior part of the flank lack these pores (Lund & Poplin, 1997: fig. 6). This taxon is scored '1'.)

0 absent

1 present

**206. Small scales below dorsal fin**

(Giles et al. 2017; Argyriou et al., 2018; Latimer & Giles, 2018; Wilson et al., 2019; Figueroa et al., 2021.)

0 absent

1 present

**207. [G 158] Lepidotrichia**

(Friedman, 2007; Brazeau, 2009; Zhu et al., 2009; Friedman & Brazeau, 2010; Davis et al., 2012; Zhu et al., 2013; Brazeau & Friedman, 2014; Giles et al., 2015b,c; Giles et al. 2017; Argyriou et al., 2018; Latimer & Giles, 2018; Wilson et al., 2019; Figueroa et al., 2021. )

0 absent

1 present

**208. [CH 37; G 159] Fringing fulcra**

(Patterson, 1982; Gardiner & Schaeffer, 1989; Grande & Bemis 1998; Coates, 1999; Dietze, 2000; Schultze & Cumbaa, 2001; Poyato-Ariza & Wenz, 2002; Cloutier & Arratia, 2004; Friedman & Blom, 2006; Friedman, 2007; Long et al., 2008; Swartz, 2009; Zhu et al., 2009; Choo, 2011; Xu & Gao, 2011; López-Arbarello 2012; Zhu et al., 2013; Xu et al., 2014, 2015; Bermúdez-Rochas & Poyato-Ariza 2015; Poyato-Ariza 2015; Giles et al., 2015b; Thies & Waschkewitz, 2015; Gibson 2016; Xu & Zhao, 2016; Giles et al. 2017; Cawley & Kriwet 2018; López-Arbarello & Sferco 2018; Argyriou et al., 2018; Latimer & Giles, 2018; Wilson et al., 2019; Figueroa et al., 2021.)

0 absent

1 present

**209. [G 139] Double headed hyomandibular**

(Cloutier & Ahlberg, 1996; Zhu & Schultze, 2001; Schultze & Cumbaa, 2001; Zhu et al., 2001; Zhu & Yu, 2002; Zhu et al., 2006; Friedman, 2007; Zhu et al., 2009; Friedman & Brazeau, 2010; Zhu et al., 2013; Giles et al., 2015b,c; Giles et al. 2017; López-Arbarello & Sferco 2018; Argyriou et al., 2018; Latimer & Giles, 2018; Wilson et al., 2019; Figueroa et al., 2021.)

0 absent

1 present

**210. [G 140] Perforate hyomandibula**

(Friedman, 2007; Zhu et al., 2009; Friedman & Brazeau, 2010; Xu & Gao, 2011; Zhu et al., 2013; Brazeau & Friedman, 2014; Xu et al., 2014, 2015; Giles et al., 2015b; Xu & Zhao, 2016; Giles et al. 2017; Argyriou et al., 2018; Latimer & Giles, 2018; Wilson et al., 2019; Figueroa et al., 2021. Although Long (1988: p.24) mentions the presence of a depression for the hyomandibular nerve in *Howqualepis*, it is unclear whether this perforated the hyomandibula. This taxon is conservatively coded as '?'.)

0 absent

1 present

**211. [G 141] Opercular process**

(Gardiner & Schaeffer, 1989 Poyato-Ariza & Wenz, 2002;; Giles et al., 2015b; Giles et al. 2017; Cawley & Kriwet 2018; López-Arbarello & Sferco 2018; Argyriou et al., 2018; Latimer & Giles, 2018; Wilson et al., 2019; Figueroa et al., 2021.)

0 absent

1 present

**212. [G 136] Ceratohyal**

(Gardiner et al., 2005; Xu & Gao, 2011; Giles et al., 2015b; Giles et al. 2017; Xu et al., 2014; Argyriou et al., 2018; Latimer & Giles, 2018; Wilson et al., 2019; Figueroa et al., 2021.)

0 single ossification

1 two ossifications

**213. [G 137] Anterior ossification of ceratohyal**

(Revised from Coates, 1999; Giles et al., 2015b; Giles et al. 2017; López-Arbarello & Sferco 2018; Argyriou et al., 2018; Latimer & Giles, 2018; Wilson et al., 2019; Figueroa et al., 2021. The character captures whether the ceratohyal (or the anterior ossification if an anterior and posterior ceratohyal are present) is medially constricted (hourglass-shaped) or plate-like in lateral view.)

0 no medial constriction

1 medial constriction (hourglass-shaped)

**214. [G 138] Anterior ceratohyal**

(Coates, 1999; Giles et al., 2015b; Giles et al. 2017; Argyriou et al., 2018; Latimer & Giles, 2018; Wilson et al., 2019; Figueroa et al., 2021. The groove for the afferent hyoidean artery in the ceratohyal of *Gogosardina* is visible in Choo 2009 (fig 6).)

0 no groove

1 groove for afferent hyoidean artery

**215. [G 144] Interhyal**

(Davis et al., 2012; Zhu et al., 2013; Giles et al., 2015b,c; Giles et al. 2017; Argyriou et al., 2018; Latimer & Giles, 2018; Wilson et al., 2019; Figueroa et al., 2021.)

0 absent

1 present

**216. Symplectic**

(Gardiner 1984; Gardiner & Schaeffer 1989; Coates 1999; Hurley et al. 2007; Poyato-Ariza 2015; Xu & Zhao, 2016; Giles et al. 2017; López-Arbarello & Sferco 2018; Argyriou et al., 2018; Latimer & Giles, 2018; Wilson et al., 2019; Figueroa et al., 2021. The general actinopterygian condition of the hyoid arch seems to comprise four ossifications: hyomandibula, ceratohyal (which may be one or two bones), hypohyal, and an intermediate bone between the hyomandibula and ceratohyal termed, variably, the interhyal or symplectic. In some actinopts (e.g. *Amia*, *Lepisosteus*, *Hiodon*, *Dorsetichthys*, *Macrosemionotus*, etc.), a second intermediate cartilage is present. The history attached to naming these terms is very complex (see Paterson 1973, Patterson 1982, Véran 1988, Gardiner et al. 1996, etc.), and we have tried here to apply a simple, consistent approach. The ossification that forms an intermediary between the hyomandibula and ceratohyal is termed the interhyal. This is primitively present (and in contact with the quadrate), and may be very reduced (e.g. *Watsonulus*, Elops), or entirely cartilaginous (e.g. *Amia*, *Lepisosteus*) in more derived actinopts. The ossification that contacts the hyomandibula (and typically the quadrate), but does not articulate with the ceratohyal, is termed the symplectic. This element may brace the quadrate, and in *Watsonulus*, *Caturus* and *Amia* additionally articulates with the lower jaw. We follow Grande (2010) in identifying the ‘symplectic’ of *Acipenser* as the posterior ceratohyal. Véran (1988) identified a second intermediate ossification in the hyoid arch in a number of ‘palaeoniscids’, which she termed a symplectic. This identification has been disputed on the basis of position (e.g. Gardiner et al. 1996). We have seen no evidence (either through visual examination or CT scanning) for a second intermediate hyoid ossification in any specimens of *Boreosomus* or *Pteronisculus*. From examination of *Coccocephalichthys*, we identify the ‘symplectic’ of Véran to be the interhyal and the ‘interhyal’ of Véran to correspond to the articular. This casts doubt on the presence of a second element, and we have therefore coded *Boreosomus*, *Pteronisculus* and *Coccocephalicthys* as ‘0’.)

0 absent

1 present

**217. Symplectic shape c68**

(Giles et al. 2017; López-Arbarello & Sferco 2018; Argyriou et al., 2018; Latimer & Giles, 2018; Wilson et al., 2019; Figueroa et al., 2021.)

0 tube/splint like

1 hatchet

2 l-shaped

**218. [G 145] Hypohyal**

(Friedman & Brazeau, 2010; Brazeau & Friedman, 2014; Giles et al., 2015b,c; Giles et al. 2017; Argyriou et al., 2018; Latimer & Giles, 2018; Wilson et al., 2019; Figueroa et al., 2021.)

0 absent

1 present

**219. [G 143] Basihyal**

(Davis et al., 2012; Zhu et al., 2013; Giles et al., 2015b,c; Giles et al. 2017; López-Arbarello & Sferco 2018; Argyriou et al., 2018; Latimer & Giles, 2018; Wilson et al., 2019; Figueroa et al., 2021.)

0 absent

1 present

**220. [G 146] Gill arches**

(Giles et al., 2015b,c; Giles et al. 2017; Argyriou et al., 2018; Latimer & Giles, 2018; Wilson et al., 2019; Figueroa et al., 2021.)

0 largely restricted to area under braincase

1 extend far posterior to braincase

**221. Number of ceratobranchials**

(Giles et al. 2017; Argyriou et al., 2018; Latimer & Giles, 2018; Wilson et al., 2019; Figueroa et al., 2021.)

0 five

1 four

**222. Number of hypobranchials**

(Grande, 2010; Xu & Wu, 2012; Xu et al., 2014, 2015; Xu & Zhao, 2016; Giles et al. 2017; López-Arbarello & Sferco 2018; Argyriou et al., 2018; Latimer & Giles, 2018; Wilson et al., 2019; Figueroa et al., 2021. )

0 three

1 four

**223. Uncinate processes on epibranchials**

(Coates, 1999; Xu & Gao, 2011; Xu et al., 2014, 2015; Poyato-Ariza 2015; Xu & Zhao, 2016; Giles et al. 2017; López-Arbarello & Sferco 2018; Argyriou et al., 2018; Latimer & Giles, 2018; Wilson et al., 2019; Figueroa et al., 2021. An uncinate process is a dorsally-directed extension on the epibranchial that articulates with the pharyngobranchial skeleton. )

0 absent

1 present

**224. [G 142] Endoskeletal urohyal**

(Friedman, 2007; Friedman & Brazeau, 2010; Giles et al., 2015b,c; Poyato-Ariza 2015; Giles et al. 2017; López-Arbarello & Sferco 2018; Argyriou et al., 2018; Latimer & Giles, 2018; Wilson et al., 2019; Figueroa et al., 2021.)

0 absent

1 present

**225. Urohyal formed as a tendon bone of the sternohyoideus muscle**

(Arratia 2013; Giles et al. 2017; Argyriou et al., 2018; Latimer & Giles, 2018; Wilson et al., 2019; Figueroa et al., 2021.)

0 absent

1 present

**226. [CH 39; G 162] Presupracleithrum**

(Patterson, 1982; Gardiner, 1984; Gardiner & Schaeffer, 1989; Taverne, 1997; Lund, 2000; Schultze & Cumbaa, 2001; Zhu & Schultze, 2001; Zhu et al., 2001; Lund & Poplin, 2002; Zhu & Yu, 2002; Cloutier & Arratia, 2004; Gardiner et al., 2005; Friedman & Blom, 2006; Zhu et al., 2006; Friedman, 2007; Long et al., 2008; Swartz, 2009; Zhu et al., 2009; Choo, 2011; Xu & Gao, 2011; Zhu et al., 2013; Xu et al., 2014; Giles et al., 2015b; Giles et al. 2017; López-Arbarello & Sferco 2018; Argyriou et al., 2018; Latimer & Giles, 2018; Wilson et al., 2019; Figueroa et al., 2021. An elongate bone termed the 'anocleithrum' is variably present in *Wendyichthys* (Lund & Poplin, 1997) in the position occupied by the presupracleithrum in other taxa. We regard this as a positional homologue, and code the taxon '0/1' to reflect this polymorphism. Coded as '?' in *C. trailli* following arguments in Friedman & Blom (2006).)

0 absent

1 present

**227. Presupracleithrum**

(Xu et al. 2014; Giles et al. 2017; Argyriou et al., 2018; Latimer & Giles, 2018; Wilson et al., 2019; Figueroa et al., 2021.)

0 single

1 multiple

**228. [G 160] Dorsal margin of cleithrum**

(Cloutier & Ahlberg, 1996; Schultze & Cumbaa, 2001; Zhu & Schultze, 2001; Zhu et al., 2001; Zhu & Yu, 2002; Cloutier & Arratia, 2004; Zhu et al., 2006; Friedman, 2007; Zhu et al., 2009; Giles et al., 2015b,c; Giles et al. 2017; Argyriou et al., 2018; Latimer & Giles, 2018; Wilson et al., 2019; Figueroa et al., 2021.)

0 pointed

1 broad and rounded

**229. Medial wing on cleithrum**

(Cavin & Suteethorn 2006; Poyato-Ariza 2015; Giles et al. 2017; López-Arbarello & Sferco 2018; Argyriou et al., 2018; Latimer & Giles, 2018; Wilson et al., 2019; Figueroa et al., 2021.)

0 absent

1 present

**230. [G 161] Anocleithrum**

(Gardiner & Schaeffer, 1989; Lund et al., 1995; Cloutier & Ahlberg, 1996; Dietze, 2000; Poplin & Lund, 2000; Schultze & Cumbaa, 2001; Zhu & Schultze, 2001; Zhu et al., 2001; Zhu & Yu, 2002; Cloutier & Arratia, 2004; Zhu et al., 2006; Friedman, 2007; Zhu et al., 2009; Zhu et al., 2013; Giles et al., 2015b; Giles et al. 2017; López-Arbarello & Sferco 2018; Argyriou et al., 2018; Latimer & Giles, 2018; Wilson et al., 2019; Figueroa et al., 2021.)

0 bone developed as postcleithrum

1 bone developed as anocleithrum sensu stricto

2 bone absent

**231. Clavicle**

(Patterson 1973; Olsen & McCune 1991; Coates 1999; Xu & Gao 2011; Xu et al. 2014; Poyato-Ariza 2015; Xu & Zhao, 2016; Giles et al. 2017; López-Arbarello & Sferco 2018; Argyriou et al., 2018; Latimer & Giles, 2018; Wilson et al., 2019; Figueroa et al., 2021.)

0 present as a broad plate

1 much reduced or absent

**232. Serrated organ**

(Arratia 2013; Giles et al. 2017; López-Arbarello & Sferco 2018; Argyriou et al., 2018; Latimer & Giles, 2018; Wilson et al., 2019; Figueroa et al., 2021. The serrated organ (or appendage) is a small, elongate element, typically ornamented with serrated ridges, present near the anterior margin of the cleithrum. )

0 absent

1 present

**233. Interclavicle**

(Cloutier & Arratia, 2004; Giles et al. 2017; Xu et al., 2014; Argyriou et al., 2018; Latimer & Giles, 2018; Wilson et al., 2019; Figueroa et al., 2021. )

0 present

1 absent

**234. [CH 40; G 167] Triradiate scapulocoracoid**

(Olsen & McCune 1991; Zhu & Schultze, 2001; Zhu et al., 2001; Zhu & Yu, 2002; Zhu et al., 2006; Friedman, 2007; Zhu et al., 2009; Xu & Gao, 2011; Zhu et al., 2013; Xu et al., 2014; Giles et al., 2015b; Giles et al. 2017; Argyriou et al., 2018; Latimer & Giles, 2018; Wilson et al., 2019; Figueroa et al., 2021.)

0 absent

1 present

**235. [G 163] Perforate propterygium**

(Patterson, 1982; Gardiner, 1984; Gardiner & Schaeffer, 1989; Rosen, 1989; Taverne, 1997; Coates, 1999; Zhu & Schultze, 2001; Zhu et al., 2001; Zhu & Yu, 2002; Zhu et al., 2006; Brazeau, 2009; Zhu et al., 2009; Friedman & Brazeau, 2010; Xu & Gao, 2011; Davis et al., 2012; Zhu et al., 2013; Xu et al., 2014; Giles et al., 2015b,c; Argyriou et al., 2018; Latimer & Giles, 2018; Wilson et al., 2019; Figueroa et al., 2021.)

0 absent

1 present

**236. [CH 41; G 164] Anterior rays embrace propterygium**

(Patterson, 1982; Gardiner, 1984; Gardiner & Schaeffer, 1989; Taverne, 1997; Coates, 1999; Schultze & Cumbaa, 2001; Zhu & Schultze, 2001; Friedman & Blom, 2006; Long et al., 2008; Swartz, 2009; Choo, 2011; Xu & Gao, 2011; Giles et al., 2015b; Giles et al. 2017; Argyriou et al., 2018; Latimer & Giles, 2018; Wilson et al., 2019; Figueroa et al., 2021. )

0 absent

1 present

2 fused

**237. Propterygium fused to first ray**

(Giles et al. 2017; López-Arbarello & Sferco 2018; Argyriou et al., 2018; Latimer & Giles, 2018; Wilson et al., 2019; Figueroa et al., 2021.)

0 absent

1 present

**238. [CH 43; G 168] Pectoral fin endoskeleton**

(Taverne, 1997; Coates, 1999; Friedman & Blom, 2006; Long et al., 2008; Swartz, 2009; Xu & Gao, 2011; Xu et al., 2014; Giles et al., 2015b; Giles et al. 2017; Argyriou et al., 2018; Latimer & Giles, 2018; Wilson et al., 2019; Figueroa et al., 2021.)

0 extends far beyond body wall (fins lobate)

1 barely extends beyond body wall (fins not lobate)

**239. [G 166] Pectoral fin radials**

(Zhu & Yu, 2002; Friedman, 2007; Giles et al., 2015b; Giles et al. 2017. Two series of pectoral fin radials are described (but not figured) for Cheirolepis candensis (Arratia & Cloutier, 2004). Although we consider this arrangement to be unlikely, for now this taxon is coded '1'; Argyriou et al., 2018; Latimer & Giles, 2018; Wilson et al., 2019; Figueroa et al., 2021. )

0 unjointed

1 jointed

**240. [G 169] Fin articulation**

(Zhu & Schultze, 2001; Zhu et al., 2001; Zhu & Yu, 2002; Zhu et al., 2006; Friedman, 2007; Zhu et al., 2009; Friedman & Brazeau, 2010; Zhu et al., 2013; Giles et al., 2015b,c; Giles et al. 2017; Argyriou et al., 2018; Latimer & Giles, 2018; Wilson et al., 2019; Figueroa et al., 2021. )

0 monobasal

1 polybasal

**241. [CH 44; G 171] Pectoral fin-ray segmentation**

(Coates, 1999; Friedman & Blom, 2006; Long et al., 2008; Choo, 2011; Xu & Gao, 2011; Xu et al., 2014; Giles et al., 2015b; Giles et al. 2017; Argyriou et al., 2018; Latimer & Giles, 2018; Wilson et al., 2019; Figueroa et al., 2021.)

0 roughly even segmentation to fin base

1 proximal segments elongate with terminal segmentation

2 no significant segmentation on pectoral fin

3 terminal segments elongate with proximal segmentation

**242. [G 170] Pectoral fin**

(Giles et al., 2015b; Giles et al. 2017; Argyriou et al., 2018; Latimer & Giles, 2018; Wilson et al., 2019; Figueroa et al., 2021.)

0 [leaf-like]

1 (not leaf-like)

**243. [G 172] Paired fin spines**

(Zhu et al., 2001; Zhu & Yu, 2002; Friedman, 2007; Zhu et al., 2009; Brazeau, 2009; Davis et al., 2012; Zhu et al., 2013; Giles et al., 2015b,c; Giles et al. 2017; Argyriou et al., 2018; Latimer & Giles, 2018; Wilson et al., 2019; Figueroa et al., 2021.)

0 absent

1 present

**244. [CH 38; G 173] Pelvic fins**

(Friedman & Blom, 2006; Friedman, 2007; Brazeau, 2009; Choo, 2011; Davis et al., 2012; Zhu et al., 2013; Brazeau & Friedman, 2014; Giles et al., 2015b,c; Giles et al. 2017; Argyriou et al., 2018; Latimer & Giles, 2018; Wilson et al., 2019; Figueroa et al., 2021. )

0 absent

1 present

**245. [CH 45; G 174] Pelvic fin insertion**

(Gardiner & Schaeffer, 1989; Coates, 1998; Coates, 1999; Lund, 2000; Schultze & Cumbaa, 2001; Cloutier & Arratia, 2004; Friedman & Blom, 2006; Zhu et al., 2006; Long et al., 2008; Swartz, 2009; Zhu et al., 2009; Choo, 2011; Xu et al., 2014; Giles et al., 2015b; Giles et al. 2017; Argyriou et al., 2018; Latimer & Giles, 2018; Wilson et al., 2019; Figueroa et al., 2021.)

0 shorter than fin depth (short based)

1 longer than fin depth (long based)

**246. [G 177] Basal scutes on fins**

(Friedman, 2007; Zhu & Yu, 2002; López-Arbarello 2012; Bermúdez-Rochas & Poyato-Ariza 2015; Giles et al., 2015b; Thies & Waschkewitz, 2015; Gibson 2016; Giles et al. 2017; López-Arbarello & Sferco 2018; Argyriou et al., 2018; Latimer & Giles, 2018; Wilson et al., 2019; Figueroa et al., 2021.)

0 absent

1 present

**247. [CH 48; G 178] Dorsal scutes anterior to dorsal fin**

(Olsen & McCune 1991; Lund, 2000; Poplin & Lund, 2000; Poyato-Ariza & Wenz, 2002; Cloutier & Arratia, 2004; Friedman & Blom, 2006; Long et al., 2008; Swartz, 2009; Choo, 2011; Giles et al., 2015b; Thies & Waschkewitz, 2015; Gibson 2016; Giles et al. 2017; Cawley & Kriwet 2018; López-Arbarello & Sferco 2018; Argyriou et al., 2018; Latimer & Giles, 2018; Wilson et al., 2019; Figueroa et al., 2021.)

0 absent

1 few limited to region immediately anterior to fin (basal fulcra only)

2 many, extending to posterior of skull roof (complete set of dorsal ridge scales)

**248. [CH 49; G 179] Ventral scutes between hypochordal lobe of caudal fin and anal fin** (Patterson, 1982; Taverne, 1997; Friedman & Blom, 2006; Long et al., 2008; Choo, 2011; Giles et al., 2015b; Giles et al. 2017; López-Arbarello & Sferco 2018; Argyriou et al., 2018; Latimer & Giles, 2018; Wilson et al., 2019; Figueroa et al., 2021.)

0 absent

1 present

**249. [CH 50; G 180] Ventral scutes anterior to anal fin**

(Cloutier & Arratia, 2004; Friedman & Blom, 2006; Long et al., 2008; Swartz, 2009; Choo, 2011; Giles et al., 2015b; Giles et al. 2017; Argyriou et al., 2018; Latimer & Giles, 2018; Wilson et al., 2019; Figueroa et al., 2021.)

0 absent

1 present

**250. [CH 52; G 181] Dorsal fin(s)**

(Gardiner & Schaeffer, 1989; Schultze & Cumbaa, 2001; Zhu & Schultze, 2001; Zhu et al., 2001; Zhu & Yu, 2002; Cloutier & Arratia, 2004; Friedman & Blom, 2006; Zhu et al., 2006; Friedman, 2007; Long et al., 2008; Brazeau, 2009; Swartz, 2009; Zhu et al., 2009; Choo, 2011; Davis et al., 2012; Zhu et al., 2013; Giles et al., 2015b,c; Giles et al. 2017; Argyriou et al., 2018; Latimer & Giles, 2018; Wilson et al., 2019; Figueroa et al., 2021.)

0 two

1 one

**251. [CH 51; G 182] Relative positions of anal and (second) dorsal fin**

(Poplin & Lund, 2000; Cloutier & Arratia, 2004; Friedman & Blom, 2006; Long et al., 2008; Swartz, 2009; Choo, 2011; López-Arbarello 2012; Bermúdez-Rochas & Poyato-Ariza 2015; Giles et al., 2015b; Thies & Waschkewitz, 2015; Gibson 2016; Giles et al. 2017; López-Arbarello & Sferco 2018; Argyriou et al., 2018; Latimer & Giles, 2018; Wilson et al., 2019; Figueroa et al., 2021.)

0 anal shifted anteriorly relative to dorsal

1 fins opposite one another

2 anal shifted posteriorly relative to dorsal

**252. Median fins (except caudal fin)**

(Gardiner et al., 2005; Xu & Gao, 2011; Xu et al., 2014, 2015; Xu & Zhao, 2016; Giles et al. 2017; López-Arbarello & Sferco 2018; Argyriou et al., 2018; Latimer & Giles, 2018; Wilson et al., 2019; Figueroa et al., 2021.)

0 rays more numerous than radials

1 rays andradials equal

**253. Proximal and middle radials of dorsal fin**

(Giles et al. 2017; Argyriou et al., 2018; Latimer & Giles, 2018; Wilson et al., 2019; Figueroa et al., 2021.)

0 proximal and middle radials of similar size

1 proximal radials substantially enlarged

**254. Posteriormost proximal radial of dorsal fin**

(Giles et al. 2017; Argyriou et al., 2018; Latimer & Giles, 2018; Wilson et al., 2019; Figueroa et al., 2021.)

0 enlarged plate

1 smaller than more anterior radials

**255. [CH 46; G 175] Epichordal lobe of caudal fin**

(Patterson, 1982; Cloutier & Ahlberg, 1996; Coates, 1999; Schultze & Cumbaa, 2001; Zhu & Schultze, 2001; Friedman & Blom, 2006; Long et al., 2008; Swartz, 2009; Choo, 2011; Giles et al., 2015b; Giles et al. 2017; Argyriou et al., 2018; Latimer & Giles, 2018; Wilson et al., 2019; Figueroa et al., 2021.)

0 present

1 absent

**256. [CH 47; G 176] Fulcra along dorsal ridge of caudal fin**

(Patterson, 1982; Taverne, 1997; Gardiner & Schaeffer, 1989; Gardiner et al., 2005; Friedman & Blom, 2006; Long et al., 2008; Choo, 2011; López-Arbarello 2012; Bermúdez-Rochas & Poyato-Ariza 2015; Giles et al., 2015b; Giles et al. 2017; López-Arbarello & Sferco 2018; Argyriou et al., 2018; Latimer & Giles, 2018; Wilson et al., 2019; Figueroa et al., 2021. )

0 absent

1 present

**257. Caudal fin geometry**

(Modified from Gardiner et al., 2005; Giles et al. 2017; Argyriou et al., 2018; Latimer & Giles, 2018; Wilson et al., 2019; Figueroa et al., 2021. A long chordal lobe is considered to be present when the notochord reaches the posterior margin of the caudal fin.)

0 long chordal lobe

1 short chordal lobe

**258. Posterior margin of caudal fin**

(Grande & Bemis 1998; Xu & Gao 2011; Xu et al. 2014; Xu & Zhao, 2016; Giles et al. 2017; López-Arbarello & Sferco 2018; Argyriou et al., 2018; Latimer & Giles, 2018; Wilson et al., 2019; Figueroa et al., 2021.)

0 forked

1 unforked

**259. Diplospondyly in mid-caudal region**

(Grande & Bemis 1998; Arratia 2013; Poyato-Ariza 2015; Giles et al. 2017; López-Arbarello & Sferco 2018; Argyriou et al., 2018; Latimer & Giles, 2018; Wilson et al., 2019; Figueroa et al., 2021.)

0 absent

1 present

**260. Median neural spines in caudal region**

(Coates 1999; Hurley et al. 2007; Xu et al. 2014; Giles et al. 2017; López-Arbarello & Sferco 2018; Argyriou et al., 2018; Latimer & Giles, 2018; Wilson et al., 2019; Figueroa et al., 2021.)

0 absent

1 present

**261. Uroneural**

(Pinna, 1996; Hurley et al., 2007; Xu & Wu, 2012, Xu et al., 2014; Xu & Zhao, 2016; Giles et al. 2017; López-Arbarello & Sferco 2018; Argyriou et al., 2018; Latimer & Giles, 2018; Wilson et al., 2019; Figueroa et al., 2021.)

0 absent

1 present

**262. Division of hypurals into dorsal and ventral groups**

(Pinna, 1996, Xu & Wu, 2012, Xu et al., 2014; Xu & Zhao, 2016; Giles et al. 2017; Argyriou et al., 2018; Latimer & Giles, 2018; Wilson et al., 2019; Figueroa et al., 2021.)

0 absent

1 present

**263. Number of caudal lepidotrichs borne per hypural**

(Giles et al. 2017; López-Arbarello & Sferco 2018; Argyriou et al., 2018; Latimer & Giles, 2018; Wilson et al., 2019; Figueroa et al., 2021.)

0 multiple	(single)

1 single

**264. Opistocoelous vertebrae**

(Wiley, 1976; López-Arbarello 2012; Bermúdez-Rochas & Poyato-Ariza 2015; Thies & Waschkewitz, 2015; Poyato-Ariza 2015; Gibson 2016; Giles et al. 2017; López-Arbarello & Sferco 2018; Argyriou et al., 2018; Latimer & Giles, 2018; Wilson et al., 2019; Figueroa et al., 2021.)

0 absent

1 present

**265. Ossified ribs**

(Giles et al. 2017; Argyriou et al., 2018; Latimer & Giles, 2018; Wilson et al., 2019; Figueroa et al., 2021.)

0 present

1 absent

**266. Antorbital deeper than anteriormost infraorbital**

(L?pez-Arbarello 2012; Thies & Waschkewitz, 2015; Gibson 2016; Lóiiiiiiiiipez-Arbarello & Sferco 2018; Latimer & Giles, 2018.)

0 absent

1 present

**267. Infraorbitals fragmented**

(Latimer & Giles, 2018. The infraorbitals normally form as a single row partially surrounding the orbit. In some taxa, they may be fragmented into multiple irregular ossifications.)

0 absent

1 present

**268. Suborbitals fragmented**

(Grande & Bemis 1998; López-Arbarello 2011; López-Arbarello & Sferco 2018; Latimer & Giles, 2018. The suborbitals normally form as a single row partially surrounding the orbit, posterior and/or ventral to the infraorbitals. In some taxa, they may be fragmented into multiple irregular ossifications.)

0 absent

1 present

**269. Suborbitals extend ventral to orbit**

(López-Arbarello 2011; López-Arbarello & Sferco 2018; Latimer & Giles, 2018.)

0 absent

1 present

**270. Coronoids contribute to lateral dentition field**

(Latimer & Giles, 2018. Coronoids are normally restricted to the medial (lingual) surface of the jaw. In some dapediids, they may extend laterally to contribute to the labial tooth row.)

0 absent

1 present

**271. Jaw articulation ventral to orbit**

(López-Arbarello 2012; Bermúdez-Rochas & Poyato-Ariza 2015; Poyato-Ariza 2015; Thies & Waschkewitz, 2015; Gibson 2016; López-Arbarello & Sferco 2018; Latimer & Giles, 2018.)

0 absent

1 present

**272. Parasphenoid wings around basiocciput**

(Latimer & Giles, 2018. The parasphenoid is normally restricted to covering the ventral face of the braincase. In some taxa, extensions may extend dorsally and laterally to cloak the basiocciput.)

0 absent

1 present

**273. Dorsal extension of parasphenoid between orbits**

(Latimer & Giles, 2018. The parasphenoid extends dorsally between the orbits, contributing to the interorbital septum, in some taxa.)

0 absent

1 present

**274. Ligament pit on posterior face of braincase**

(Latimer & Giles, 2018. A deep ligament pit, dorsal to the foramen magnum, is found on the posterior face of the braincase in some taxa.)

0 absent

1 present

**275. 'Hem-like' median fins**

(Gibson 2016; Latimer & Giles, 2018.)

0 absent

1 present

**276. Anal fin insertion**

(Latimer & Giles, 2018. )

0 short

1 long

**277. Denticulate ridge scales**

(Latimer & Giles, 2018.)

0 absent

1 present

**278. Ossified centra**

(Poyato-Ariza & Wenz, 2002; Cawley & Kriwet 2018; Latimer & Giles, 2018.)

0 absent

1 present

**279. Supratemporal commissure in parietal**

(Poyato-Ariza & Wenz, 2002; Thies & Waschkewitz, 2015; Gibson 2016; Cawley & Kriwet 2018; López-Arbarello & Sferco 2018; Latimer & Giles, 2018. The supratemporal commissure is normally carried by the extrascapulars, but is sometimes borne in the parietal.)

0 absent

1 present

**280. Enlarged vomerine teeth**

(Poyato-Ariza & Wenz, 2002; Cawley & Kriwet 2018; López-Arbarello & Sferco 2018; Latimer & Giles, 2018.)

0 absent

1 present

**281. Enlarged prearticular teeth**

(Poyato-Ariza & Wenz, 2002; Cawley & Kriwet 2018; Latimer & Giles, 2018.)

0 absent

1 present

**282. Prearticular symphysis**

(Nursall 2010.; Latimer & Giles, 2018)

0 absent

1 present

**283. Dermal supraoccipital**

(Poyato-Ariza & Wenz, 2002; Cawley & Kriwet 2018; Latimer & Giles, 2018. A median bone, typically referred to as a dermal supraoccipital, is present between the parietals in some taxa.)

0 absent

1 present

**284. Preoperculum taller than operculum**

(Poyato-Ariza & Wenz, 2002; Cawley & Kriwet 2018; Latimer & Giles, 2018.)

0 shorter

1 taller

**285. Preoperculum wider than operculum**

(Poyato-Ariza & Wenz, 2002; Cawley & Kriwet 2018; Latimer & Giles, 2018.)

0 narrower

1 wider

**286. Branchiostegals contact next in sequence**

(Poyato-Ariza & Wenz, 2002; Cawley & Kriwet 2018; Latimer & Giles, 2018.)

0 absent

1 present

**287. Dorsal lateral line**

(Latimer & Giles, 2018.)

0 absent

1 present

**288. Parasphenoid inflected ventrally below orbit**

(Nursall 2010; Latimer & Giles, 2018.)

0 absent

1 present

**289. Sagittal flange on neural spine**

(Poyato-Ariza & Wenz, 2002; Cawley & Kriwet 2018; Latimer & Giles, 2018.)

0 absent

1 present

**290. sagittal flange on haemal spine**

(Poyato-Ariza & Wenz, 2002; Cawley & Kriwet 2018; Latimer & Giles, 2018.)

0 absent

1 present

**291. Pleural ribs alate**

(Nursall 2010; Latimer & Giles, 2018.)

0 absent

1 present

**292. Concavity on the ventral margin of the dentary forming the border of the lower jaw** (Figueroa et al. 2021; Latimer & Giles, 2018.)

0 absent

1 present

| **Taxon** |   Source |
| --- | --- |
| *Acanthodes bronni* | Miles, 1973; Davis et al., 2012. |
| *Acipenser brevirostrum* | Nieuwenhuys, 1982; Hilton et al., 2011. |
| *Aesopichthys erinaceus* | Poplin & Lund, 2000. |
| *Aetholepis mirablis* | Woodward 1895. |
| *Amia calva* | Grande & Bemis, 1998; Sallan, 2014. |
| *Amphicentrum granulosum* | Traquair, 1875; Bradley-Dyne, 1939. |
| *Arduafrons prominoris* | Nursall 1999. |
| *Atractosteus spatula* | Grande, 2010. |
| *Austelliscus ferox* | Figueroa et al., 2021. |
| *Avonichthys manskyi* | Wilson et al. 2018. |
| *Cuneognathus gardineri* | Friedman & Blom, 2006. |
| *Dipteronotus ornatus* | Nielsen, 1949. |
| *Beagiascus pulcherrimus* |   Mickle et al., 2009.   |
| *Beishanichthys brevicaudalis* | Xu & Gao, 2011. |
| *Birgeria groenlandica* |   Nielsen, 1949. |
| *Bobasatrania groenlandica* |   Stensiö, 1932.   |
| *Boreosomus piveteaui* | Nielsen, 1942. |
| *Brembodus ridens* |   Tintori 1981.   |
| *Caturus furcatus* | Patterson, 1975; Lambers, 1994; Grande & Bemis, 1998. |
| *Cheirolepis canadensis* |   Arratia & Cloutier, 1996; Arratia, 2009. |
| *Cheirolepis schultzei* |   Arratia & Cloutier, 2004.   |
| *Cheirolepis trailli* | Pearson & Westoll, 1979; Giles et al., 2015a. |
| *Chondrosteus acipenseroides* |   Hilton & Forey 2009; Hilton et al., 2011. |
| *Cladodoides wildungensis* |   Maisey, 2005. |
| *Coccocephalichthys wildi* |   Poplin, 1974; Poplin & Véran, 1996. |
| *Cosmoptychius striatus* |   Schaeffer, 1971; Coates, 1999.   |
| *Cyranorhis bergeraci* | Lund et al., 1997. |
| *Dandya ovalis* |   Tintori 1983.   |
| *Dapedium caelatum* | Thies & Hauff, 2008; Thies & Herzog, 1999. |
| *Dapedium punctatum* |   Thies & Herzog, 1999; Thies & Hauff, 2011; Thies & Waschewitz, 2015. |
| *Dapedium noricum* |   Tintori 1983.   |
| *Dapedium pholidotum* | Patterson, 1975; Thies & Herzog, 1999; Thies et al., 2015. |
| *Dapedium stollorum* | Thies & Herzog, 1999; Thies & Hauff, 2011; Thies & Waschewitz, 2015. |
| *Dapedium* sp. (Lias) | Rayner 1948; Patterson 1975. |
| *Dialipina salguerioensis* | Schultze, 1968; Schultze & Cumbaa, 2001. |
| *Diplocercides kayseri* | Stensiö, 1922a,b; Forey, 1996. |
| *Dipteronotus ornatus* | Bürgin, 1992. |
| *Discoserra pectinodon* | Lund, 2000; Hurley et al., 2007.   |
| *Donnrosenia schaefferi* | Long et al., 2008.   |
| *Dorsetichthys bechei* | Patterson, 1968; Patterson, 1973; Patterson, 1975; Grande & Bemis, 1998. |
| *Ebenaqua ritchei* |   Campbell & Le Duy Phuoc, 1983. |
| *Elops hawaiensis* |   Forey, 1973. |
| *Entelognathus primordialis* |   Zhu et al., 2013.   |
| *Eomesodon liassicus* | Gardiner 1960.   |
| *Erpetoichthys calabaricus* | Claeson et al., 2007; Claeson & Hagadorn, 2008; Giles et al. 2017. |
| *Eusthenopteron foordi* |   Jarvik, 1980.   |
| *Evenkia eunoptera* | Berg, 1942; Selenzneva, 1985; Sytchevskaya, 1999. |
| *Fouldenia ischiptera* |   Sallan & Coates, 2013. |
| *Fukangichthys longidorsalis* |   Xu et al., 2014. |
| *Gibbodon cenensis* |   Tintori 1981.   |
| *Glyptolepis groenlandica* | Jarvik, 1972; Jarvik, 1980.   |
| *Gogonasus andrewsae* | Long et al., 1997; Long et al., 2006; Holland, 2014. |
| *Gogosardina coatesi* |   Choo et al., 2009.   |
| *Guiyu oneiros* | Zhu et al., 2009; Qiao & Zhu, 2010.   |
| *Hemicalypterus weiri* | Schaeffer, 1967; Gibson 2013, 2016. |
| *Heterostrophus phillipsi* |   Woodward 1928, 1929. |
| *Hiodon alosoides* |   Hilton, 2002. |
| *Howqualepis rostridens* | Long, 1988; Choo, 2009. |
| *Hulettia americana* | Schaeffer & Patterson, 1984. |
| *Ichthyokentema purbeckensis* | Griffith & Patterson, 1963. |
| *Kalops monophyrum* | Poplin & Lund 2000. |
| *Kansasiella eatoni* | Poplin, 1974. |
| *Kentuckia deani* | Rayner, 1951; Giles & Friedman, 2014. |
| *Kentuckia hlavini* | Dunkle, 1964; Feldman, 1996. |
| *Krasnoyarichthys jesseni* | Prokofiev, 2002. |
| *Lambeia pectinatus* | Mickle 2017. |
| *Lawrenciella schaefferi* | Poplin, 1984; Hamel & Poplin, 2008. |
| *Lepisosteus osseus* |   Balfour & Parker, 1882; Mathiesen & Popper, 1987; Grande, 2010. |
| *Leptolepis bronni* |   Rayner, 1937.   |
| *“Ligulalepis”* | Basden et al., 2000; Basden & Young, 2001; Clement et al., 2018. |
| *Limnomis delayni* | Daeschler, 2000. |
| *Luederia kempi* |   Schaeffer & Dalquest, 1978. |
| *Luganoia lepidosteoides* |   Bürgin, 1992. |
| *Macrepistius arenatus* |   Schaeffer, 1960; Schaeffer, 1971.   |
| *Macrosemimimus lennieri* | Jain & Robinson, 1963; Wenz, 1967; Patterson, 1875; Schröder et al., 2012. |
| *Macrosemius rostratus* |   Bartram, 1977.   |
| *Meemannia eos* | Zhu et al., 2006, 2010; Lu et al. 2016. |
| *Melanecta anneae* |   Coates, 1998. |
| *Mesturus* sp. | Nursall 1999 |
| *Mesturus verrucosus* | Nursall 1999 |
| *Miguashaia bureaui* | Cloutier, 1996. |
| *Mimipiscis bartrami* | Gardiner, 1984; Choo, 2011. |
| *Mimipiscis toombsi* | Gardiner, 1984; Choo, 2011; Giles & Friedman, 2014. |
| *Moythomasia lineata* | Jessen, 1968; Choo 2015. |
| *Moythomasia nitida* | Jessen, 1968; Choo 2015. |
| *Moythomasia perforata* | Gross, 1942. |
| *Moythomasia durgaringa* | Gardiner, 1984; Long & Trinajstic, 2010; Choo 2015. |
| *Novogonatodus kasantsevae* | Long, 1998; Holland et al., 2007. |
| *Obaichthys decoratus* | Grande, 2010. |
| *Onychodus jandemarrai* | Andrews et al., 2006. |
| *Osorioichthys marginis* | Taverne, 1997. |
| *Osteolepis macrolepidotus* | Jarvik 1948. |
| *Ozarcus mapesae* | Pradel et al., 2014. |
| *Paradapedium egertoni* | Jain 1973. |
| *Peltopleurus lissocephalus* | Bürgin, 1992. |
| *Platysomus superbus* | Moy-Thomas & Bradley-Dyne, 1938. |
| *Polypterus bichir* | Allis 1922; Jollie 1984; Bjerring 1991; Bartsch & Gemballa 1992; Bartsch et al. 1997; Grande 2010. |
| *Porolepis sp.* | Jarvik, 1972; Jarvik, 1980. |
| *Propterus elongatus* | Bartram, 1977. |
| *Psarolepis romeri* | Yu, 1998; Zhu et al., 1999; Zhu & Yu, 2009; Qu et al., 2013. |
| *Pteronisculus stensioi* | Nielsen 1942; Coates, 1998. |
| *Raynerius splendens* | Giles et al., 2015b. |
| *Sargodon tomicus* | Tintori 1983. |
| *Saurichthys madagascarensis* | Kogan et al., 2016. |
| *Scanilepis dubia* | Aldinger, 1937; Lehman, 1979. |
| *Scopulipiscis saxciput* | Giles & Latimer, 2018. |
| *Semionotus elegans* | Olsen & McCune, 1991. |
| *Stegotrachelus finlayi* | Gardiner, 1963; Swartz, 2009. |
| *Styloichthys changae* | Zhu & Yu, 2002; Zhu et al., 2006; Friedman, 2007. |
| *Styracopterus fulcratus* | Sallan & Coates, 2013. |
| *Tanaocrossus kalliokoskii* | Schaeffer, 1967; Schaeffer & Donald, 1978. |
| *Tegeolepis clarki* | Gardiner, 1963; Dunkle & Schaeffer, 1973. |
| *Tetragonolepis oldhami* | Jain 1973. |
| *Tetragonolepis semicincta* | Thies, 1991. |
| *Trawdenia planti* | Coates, 1999; Coates & Tietjen 2018. |
| *Venusichthys comptus* | Xu & Zhao, 2016. |
| *Watsonulus eugnathoides* | Olsen, 1984; Grande & Bemis, 1998. |
| *Wendyichthys dicksoni* | Lund & Poplin, 1997. |
| *Woodichthys bearsdeni* | Coates, 1998. |

**Fig. S1.** Geological context of MCZ VPF 5114 [new taxon name redacted from preprint]. (A) Map of Pennsylvania showing location; inset shows geological map for the area of Warren County (based on (36)) and simplified geological column (based on (68)). City of Warren indicated by solid bounding lines. (B) External photograph of MCZ VPF 5114 [new taxon name redacted from preprint]showing right lateral flank. Colors in stratigraphic column relate to geological map: latest Devonian and post-Devonian strata condensed into one color. Scale bar in (A): 10 km, in (B): 10 mm. Photos copyright President and Fellows of Harvard College.

**Figure S2.** External photographs of MCZ VPF 5114 [new taxon name redacted from preprint]. (A) Right lateral flank. (B) Left lateral flank. (C) Ventral view showing entire specimen, and (D) closeup of skull and pectoral region. (E) Right anal fin. (F) Scales from right lateral flank Scale bar in (A)-(C): 10 mm, in (D)-(F): 5mm. Photos copyright President and Fellows of Harvard College.

**Fig. S3.** Anatomy of MCZ VPF 5114 [new taxon name redacted from preprint]*.* Interpretive drawing of (A) right lateral view of specimen; (B) medial view of right upper and lower jaws; (C) medial view of left shoulder girdle and fin, and (D) left lateral view of gill skeleton, hyoid arch and braincase. Colour coding of skeleton: blue, cheek and jaw; purple, skull roof and sclerotic ossicle; pink, braincase; dark green, hyomandibula; light green, operculogular system; turquoise, shoulder girdle; yellow, gill skeleton. Scale bar in (A): 10 mm, in (B)-(D): 5 mm. Abbreviations: ang, angular; art, articular; asp, ascending process of parasphenoid; av, accessory vomer; br, branchiostegal; cb, ceratobranchial; chy, ceratohyal; clav, clavicle; dent, dentary; dmpt, dermatopterygoid; dsph, dermosphenotic; ecpt, ectopterygoid; eb, epibranchial; enpt, entopteryygoid; exo, exoccipital; hmd, hyomandibula; ?ic, possible intercalar; icl, interclavicle; ioc, infraorbital canal; it, intertemporal; jug, jugal; jugc, jugal canal; lep, lepidotrichia; ll, lateral line; mc, mandibular canal; mx, maxilla; op, operculum; part, prearticular and coronoid; popc, preopercular canal; popl, preopercular pit lines; ?pmx, possible premaxilla; psp, parasphenoid; qu, quadrate; rad, radials; sc, scale; scl, sclerotic ossicle; scpc, scapulocoracoid; soc, supraorbital canal; sop, suboperculum; sorb, suborbitals; sph, sphenotic ossification; st, supratemporal; t, teeth; tp, toothplate; unc, uncinate process.

Fig. S4. External anatomy of MCZ VPF 5114 [new taxon name redacted from preprint]. Renders of specimen in (A) right lateral, (B) left lateral, (C) dorsal, (D) anterior, and (E) ventral view; and an isolated scale (F). Colour coding of skeleton: blue, cheek and jaw; purple, skull roof and sclerotic ossicle; pink, braincase; dark green, hyomandibula; light green, operculogular system; turquoise, shoulder girdle; yellow, gill skeleton. Scale bar in (A)-(C),(E): 5 mm, in (D): 2 mm, in(F):1 mm. Abbreviations: ang, angular; antd, anterodorsal process; art, articular; asc, ascending process of the parasphenoid; av, accessory vomer; br, branchiostegal; cb, ceratobranchial; chy, ceratohyal; clav, clavicle; clth, cleithrum; dent, dentary; dsph, dermosphenotic; ecpt, ectopterygoid; eb, epibranchial; enpt, entopterygoid; exsc, extrascapular; fr, frontal; fu, fused lepidotrichia; hb, hypobranchial; hmd, hyomandibula; ioc, infraorbital canal; ?ic, possible intercalar; icl, interclavicle; it, intertemporal; jgl, jugal canal; jug, jugal; lac, lacrimal; lep, lepidotrichia; ll, lateral line; mc, mandibular canal; mpl, middle pit line; mx, maxilla; op, operculum; ?pmx, possible premaxilla; pa, parietal; part, prearticular and coronoids; peg, scale peg; pi, pineal foramen; pop, preoperculum; popl, preopercular pit line; ppl, posterior pit line; pscl, presupracleithrum; psp, parasphenoid; pt, posttemporal; qu, quadrate; rad, radial; sc, scale; scl, sclerotic ossicle; sclth, supracleithrum; soc, supraorbital canal; sop, suboperculum; sorb, suborbital; sph, sphenotic ossification; st, supratemporal; unc, uncinate process.

Fig. S5. Renders of MCZ VPF 5114 [new taxon name redacted from preprint]. (A) Orbit in right lateral view, and with jugal and temporal bones removed (B). (C) Possible premaxilla. (D) Skull roofing bones in visceral view. (E) Right lower jaw in dorsal view. (F) Right maxilla, dentary and postdentary in medial view. (G) Left lower jaw in medial view. Colour coding of skeleton: grey, bone; red, sensory canal. Scale bar in all: 2 mm. Abbreviations: add, adductor fossa; ang, angular; art, articular; dent, dentary; dsph, dermosphenotic; exsc, extrascapular; fr, frontal; it, intertemporal; ioc, infraorbital canal; jug, jugal; la, lacrimal; ll, lateral line; mc, mandibular canal; mx, maxillary shelf; na, nasal; ov, maxilla overlap of dentary; ?pmx, premaxilla; part, prearticular plus coronoids; pi, pineal foramen; pt, posttemporal; qu, quadrate; ram, ramifying tubules of infraorbital canal; ri, ridge; pa, parietal; scl, sclerotic ossicle; sh, maxillary shelf; soc, supraorbital canal; sorb, suborbitals; st, supratemporal; t, tooth; tp, toothplate.

Fig. S6. Renders of braincase, hyoid arch and gill skeleton of MCZ VPF 5114 [new taxon name redacted from preprint]. (A) Braincase in right lateral view. (B) Braincase in ventral view. (C) Hyoid arch, braincase and gill skeleton in right lateral view. (D) Hyoid arch in right lateral view. (E) Gill skeleton in dorsal view. (F) Gill skeleton in ventral view. Scale bar in all: 5 mm. Abbreviations: ahy, afferent hyoid artery; aoc, aortic canal; asp, ascending process of the parasphenoid; av, accessory vomer; bb, basibranchial; bhc, buccohypophysial s; boc, basioccipital; cb, ceratobranchial; chy, ceratohyal; dhy, dermohyal; d.bpt, dermal basipterygoid process; eb, epibranchial; epi.a, foramen for epibranchial artery; exo, exoccipital; hmd, hyomandibula; hb, hyobranchial; ?ic, possible intercalar; pb, pharyngobranchial; psp, parasphenoid; sph, sphenotic ossification; spig; spiracular groove.

Fig. S7. Renders of shoulder and fin of MCZ VPF 5114 [new taxon name redacted from preprint]. (A) Shoulder girdle and pectoral fin in left lateral view, and closeup with dermal girdle removed (B). (C) Shoulder girdle and pectoral fin in medial view, and with radials and lepidotrichia removed (D). (E) Shoulder girdle and pectoral fin in posterior view, and with radials and lepidotrichia removed (F). Radials in (G) dorsal and (H) ventral view. Scale bar in (A)-(F): 2 mm, in (G)-(H): 1mm. Abbreviations: clav, clavicle; clth, cleithrum; fu, fused lepidotrichia; gl, glenoid fossa; icl, interclavicle; lep, lepidotrichia; mptg, metapterygium; pr.rad, preaxial radial; ptg, propterygium; ptg.ca, propterygial canal; rad, radial; scps.hz, horizontal plate of scapulocoracoid; scpc.pl, scapulocoracoid plates.

Fig. S8. Results of parsimony analysis. (A) Strict consensus of the 20,000 trees (1616 steps) for 121 taxa and 292 equally weighted characters. Digits above nodes indicate Bremer decay indices above 1. New taxon shown in bold. (B) Agreement subtree. New taxon shown in bold.

Fig. S9. Results of Bayesian analysis. 50% majority rule tree for 121 taxa and 292 equally weighted characters. Digits at nodes indicate posterior probability support. New taxon shown in bold.

Fig. S10. Results of fossil birth-death analysis. Blue bars associated with nodes indicate 95% highest posterior density for age estimates.

Dataset S1 (separate file). Zip file containing files necessary to perform parsimony analysis in TNT (morphological data matrix, TNT script, constraint tree) as well as a consensus tree file.

Dataset S2 (separate file). Zip file morphological data matrix formatted for Bayesian analysis, as well as a consensus tree file.

Dataset S3 (separate file). Excel file containing data for timescaling and lower jaw analyses. Sheet 1 contains taxon list, data source, standard length, law length, age data locality and stratigraphy. NA indicates that lower jaw length or that the taxon was not included in our analysis. Color coding: red = inapplicable; orange = measured from literature; green = measured from specimen. Sheet 2 contains taxonomic notes. Sheet 3 contains average jaw and standard lengths for taxa in which both can be measured.

Dataset S4 (separate file). Zip file containing MrBayes script for fossil birth death analysis and consensus file.

Dataset S5 (separate file). Large format PDF of a single most parsimonious tree showing all character optimisations.
