## Supplementary figures and images for "High lineage survivorship across the end-Devonian Mass Extinction suggested by a remarkable new Late Devonian actinopterygian"

### Dataset S5 optimisation_tree.pdf

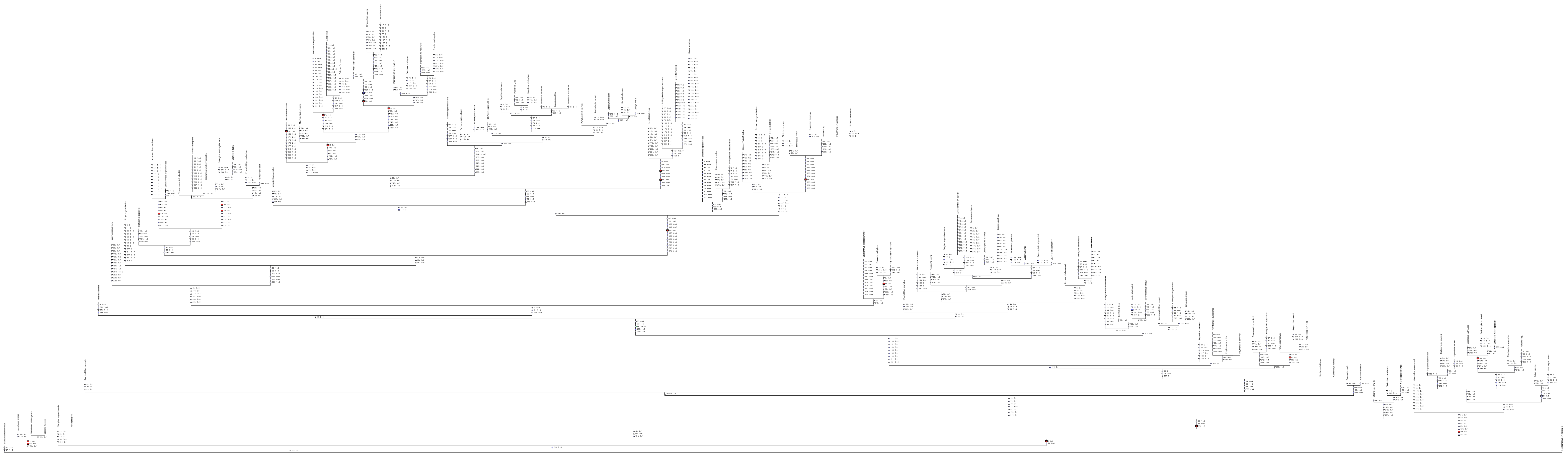
